# Safeguarding *Drosophila* female germ cell identity depends on an H3K9me3 mini domain guided by a ZAD zinc finger protein

**DOI:** 10.1101/2022.09.04.506543

**Authors:** Laura Shapiro-Kulnane, Micah Selengut, Helen K. Salz

## Abstract

H3K9me3-based gene silencing is a conserved strategy for securing cell fate, but the mechanisms controlling lineage-specific installation of this epigenetic mark remain unclear. In *Drosophila*, H3K9 methylation plays an essential role in securing female germ cell fate by silencing lineage inappropriate *phf7* transcription. Thus, *phf7* regulation in the female germline provides a powerful system to dissect the molecular mechanism underlying H3K9me3 deposition onto protein coding genes. Here we used genetic studies to identify the essential cis-regulatory elements, finding that the sequences required for H3K9me3 deposition are conserved across *Drosophila* species. Transposable elements are also silenced by an H3K9me3-mediated mechanism. But our finding that *phf7* regulation does not require the dedicated piRNA pathway components, *piwi, aub, rhino, panx*, and *nxf2*, indicates that the mechanisms of H3K9me3 recruitment are distinct. Lastly, we discovered that an uncharacterized member of the zinc finger associated domain (ZAD) containing C2H2 zinc finger protein family, IDENTITY CRISIS (IDC; CG4936), is necessary for H3K9me3 deposition onto *phf7*. Loss of *idc* in germ cells interferes with *phf7* transcriptional regulation and H3K9me3 deposition, resulting in ectopic PHF7 protein expression. IDC’s role is likely to be direct, as it localizes to a conserved domain within the *phf7* gene. Collectively, our findings support a model in which IDC guides sequence-specific establishment of an H3K9me3 mini domain, thereby preventing accidental female-to-male programming.

**Author Summary:** Tissue development and function relies on cells remembering their identity. A cell’s identity is defined by the genes it expresses and those it does not. Recent work has shown that genes can be silenced by trimethylation of histone H3 lysine 9 (H3K9me3) marked chromatin, and that H3K9me3-mediated gene silencing is a vital strategy for securing cell fate. But there is very little information about how the machinery responsible for H3K9 methylation finds its target genes. Here we explore this issue using the *Drosophila* female germline where a mini domain of repressive H3K9me3 chromatin secures female germ cell fate by silencing *phf7*, a gene normally expressed in male germ cells. Transposable elements are also silenced by H3K9me3 mini domains, but we find that the proteins involved in this process are not required for *phf7* silencing. Instead, we find that silencing requires a previously uncharacterized protein, we have named IDENTITY CRISIS. Our work provides evidence that IDENTITY CRISIS directs the H3K9 methylation machinery to build a mini domain at the *phf7* locus. Our results shed new light into how cells safeguard their identity by silencing cell type inappropriate genes, and more specifically how these genes are identified by the silencing machinery.

## Introduction

Gene silencing is critical to establishing and maintaining cell fates. Once made, the decision to silence a gene is fortified by the acquisition of repressive histone modifications. While cell type specific epigenetic silencing is primarily associated with tri-methylation of histone H3 lysine 27 (H3K27me3)-marked chromatin, tissue specific genes can also be repressed by H3K9 methylation [1–4]. For example, in *S. pombe*, discrete H3K9me3 domains silence meiotic genes in vegetative cells (e.g., [5,6]). In *C. elegans*, H3K9 methylation silences inappropriate cell type specific genes, including germline genes, in somatic cells (e.g., [7,8]). In the *D. melanogaster* female germline, H3K9me3 silences male germline genes [9]. In the mouse, H3K9me3 silences germline genes during early embryonic development (e.g., [10,11]). Studies carried out in mammalian tissue culture systems further identify H3K9me3-mediated gene silencing as a conserved and vital strategy for maintaining cell fates in a wide range of tissues (e.g., [12–23]). However, the molecular mechanisms controlling tissue specific installation of this epigenetic modification onto protein-coding genes are largely unknown.

The repressive H3K9me3 histone modification has well characterized roles in constitutive heterochromatin formation, and the transcriptional silencing of repetitive DNA elements such as satellite repeats and transposable elements (TEs) [24–26]. These studies identified two mechanisms of H3K9me3 recruitment. One mechanism involves small RNAs that guide localization through a complementary base pairing mechanism. In *Drosophila* and mammals, for example, the PIWI-associated small RNAs (piRNAs) guide the H3K9me3 silencing machinery to TEs [27]. A second mechanism involves sequence specific DNA binding proteins. In mammals, for example, H3K9me3 can be guided to TEs by members of the vertebrate specific Kruppel-associated box zinc finger (KRAB-ZNF) family of DNA binding proteins [28–31]. An analogous mechanism might exist in *Drosophila*, as two zinc finger proteins, KIPFERL and SMALL OVARY, were recently shown to have roles in TE silencing [32– 35]. Whether installation of this epigenetic modification onto protein-coding genes employs similar mechanisms remains unclear.

In *Drosophila*, H3K9 methylation plays an essential role in securing female germ cell fate by silencing lineage inappropriate *<u>PH</u>D finger protein <u>7</u>* (*phf7*) transcription [9,36]. Thus, *phf7* regulation in the female germline provides a powerful system to investigate how the H3K9me3 silencing mark is installed at protein-coding genes. *phf7* encodes a predicted chromatin reader, first identified in a screen for genes expressed in male but not female embryonic germ cells [37]. Curiously, *phf7* is not essential for male fertility, but it is critical that female germ cells not express the PHF7 protein. Forced expression of PHF7 activates a toxic gene expression program enriched for genes usually restricted to the male germline [36]. Even though PHF7 protein is limited to male germ cells, *phf7* is transcribed in both male and female germ cells. Sex specificity is achieved by alternative transcription start sites (TSS; **Fig 1A**). In testes, transcription from the upstream TSS (TSS1) produces a long mRNA isoform that makes protein. In ovaries, an H3K9me3 mini domain prevents the selection of TSS1. Instead, transcription initiates from the downstream TSS (TSS2) to produce a short mRNA isoform that is not translated and has no function. Of the three *Drosophila* enzymes known to methylate H3K9, only SETDB1 has a specific and nonredundant role in repressing *phf7* [9]. Germ cell specific loss of SETDB1, its binding partner ATF7IP, or the H3K9 reader HP1a produced ovarian germ cell tumors that inappropriately express the PHF7 protein. Notably, derepression results from losing the H3K9me3 mini domain. While these results establish the H3K9me3 mini domain controls *phf7* transcription, the mechanisms controlling installation of this epigenetic modification remains unclear.

**Fig 1.**
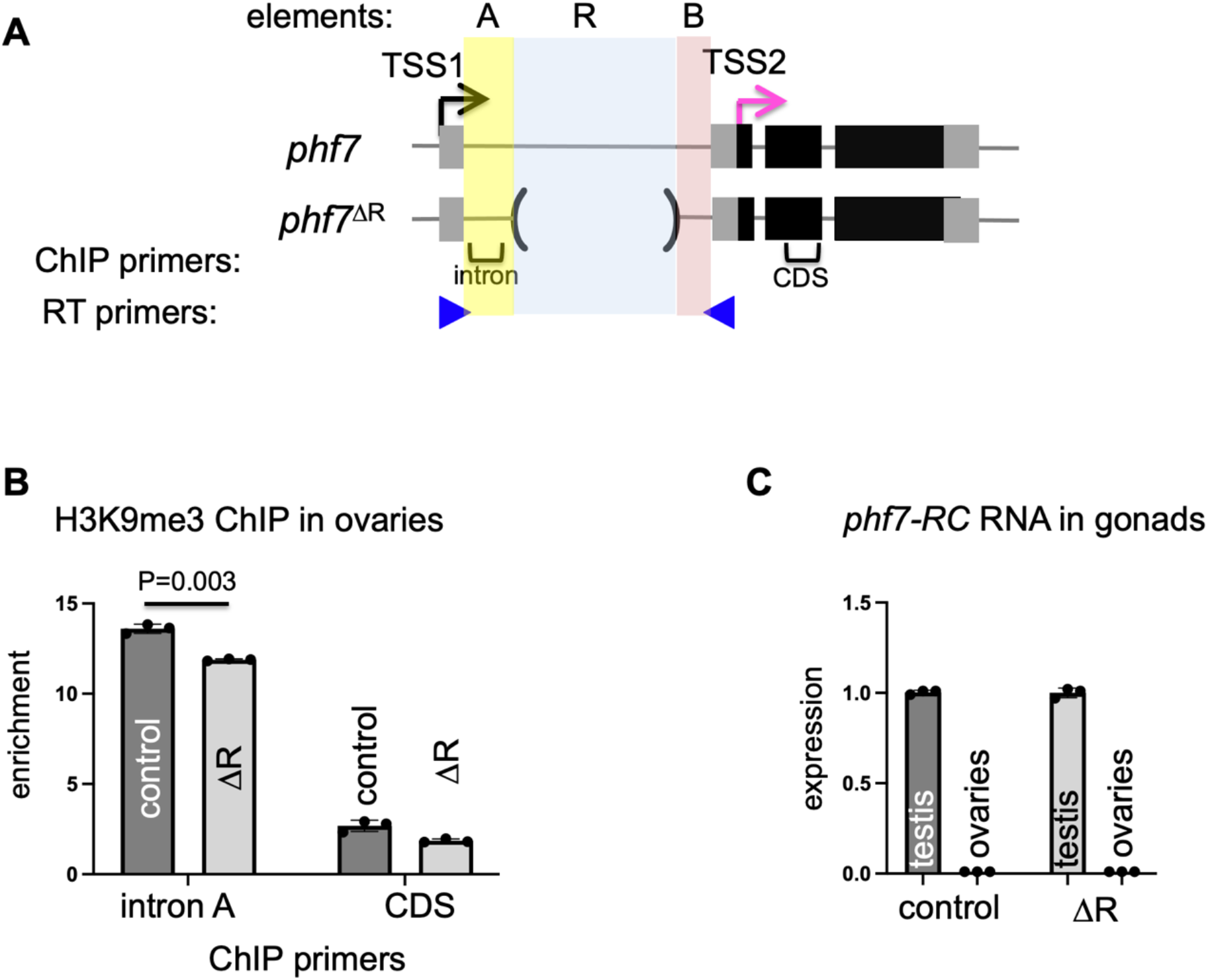
H3K9me3 deposition and sex specific transcriptional regulation are maintained in *phf7*^*ΔR*^ ovaries. (A) Diagram of the *phf7*^*+*^ and *phf7*^*ΔR*^ alleles. Exons are represented by boxes, flanking DNA, and introns by lines, black boxes are coding sequences, and grey boxes are untranslated regions (UTRs). The sequence elements A, R, and B location within the first intron are shaded in yellow, blue, and pink. In ovaries, transcription initiates from TSS2 (pink arrow), whereas an H3K9me3-mediated mechanism represses transcription from TSS1 (black arrow). Locations of primers for ChiP-qPCR and RT-qPCR are indicated by brackets & blue arrowheads. **(B)** H3K9me3 accumulation at the endogenous *phf7* locus in control (*y*^*1*^ *w*^*1*^) and mutant ovaries. ChIP-qPCR measured H3K9me3 signal at the *phf7* locus. ChIP to input enrichment is normalized to *rp49*. **(C)** Expression of the testis specific *phf7* transcript (*phf7-RC)* in control (*y*^*1*^ *w*^*1*^) and mutant ovaries. Expression is measured by RT-qPCR and is shown as the fold change relative to the testis. Expression is normalized to the total level of *phf7*. For both RT-qPCR and ChIP-qPCR experiments. Error bars represent the standard error of the mean (SD) of three biological replicates. A two-tailed Student’s t-test estimates statistical significance.

We address the mechanism of targeted H3K9me3 deposition onto *phf7* by identifying essential cis-regulatory elements and trans-acting factors. We establish that the required cis-regulatory sequences are conserved across *Drosophila* species and find that the mechanism governing *phf7* regulation is different from what has been described for piRNA-guided H3K9me3 deposition on TEs. Lastly, we discover that repression depends on the previously unknown gene, *CG4936*, that we have named *identity crisis (idc). idc* encodes a zinc finger associated domain (ZAD) containing C2H2 zinc finger protein. Notably, loss of *idc* in germ cells interferes with *phf7* repression by reducing H3K9me3 deposition. Together with the observation that IDC localizes to a conserved region within the *phf7* gene, our analysis supports a model in which IDC guides H3K9me3 installation, thereby preventing accidental female-to-male programming.

## Results

### Identification of cis-regulatory elements required for H3K9me3 deposition

H3K9me3 accumulates over a three kb region within the *phf7* gene [9]. This discrete peak covers the male TSS1, the first exon, and most of the first intron. Interestingly, the intron contains seven copies of ∼250 bp sequence not found anywhere else in the genome (element R; **Fig 1A and Fig S1**). Repetitive elements can regulate gene expression in *cis* by serving directly as H3K9me3 nucleation sites or in *trans* by encoding small RNAs that serve as guides via a base pairing mechanism. We, therefore, hypothesized that the repeats play a role in H3K9me3 deposition and silencing of TSS1. To test this idea, we used CRISPR editing to produce animals deleted for the repeats in the endogenous locus (*phf7*^*ΔR*^) (**Fig 1A**). We were surprised to discover that homozygous *phf7*^*ΔR*^ animals are viable and fertile, suggesting that the repeats may not be essential for repression. To evaluate the impact of deleting the repeats in more detail, we measured H3K9me3 accumulation in wild-type and *phf7*^*ΔR*^ homozygous ovaries using chromatin immunoprecipitation followed by quantitative PCR (ChIP-qPCR). In wild-type (control) ovaries, sequences in the intron positioned 319 bp downstream of TSS1 (within intronic element A) are significantly enriched in the H3K9me3 modification when compared to the region within the coding sequence (CDS; *p*<0.0001) (**Fig 1B**). Significant H3K9me3 enrichment within element A compared to the CDS region (*p*<0.0001) was also observed in *phf7*^*ΔR*^ homozygous ovaries. Interestingly, there was less H3K9me3 accumulation within element A in *phf7*^*ΔR*^ than in controls (*p*=0.0003), raising the possibility that sex specific transcriptional regulation might be disrupted. In testes, transcription from the upstream TSS (TSS1) produces a long mRNA isoform, called *phf7-RC*. We therefore used RT-qPCR to assay for testis specific *phf7-RC* expression levels in wild-type and *phf7*^*ΔR*^ gonads. Contrary to our expectations, we found that *phf7*^*ΔR*^ did not disrupt sex specific transcription: As in wild-type gonads, *phf7-RC* expression was detected in *phf7*^*ΔR*^ homozygous testis but not ovaries (**Fig 1C**). These results indicate that the repeats are not essential for H3K9me3-mediated repression.

To complement these studies, we used a transgenic approach to identify the sequences capable of promoting H3K9me3 to an ectopic location. For these assays we created transgenic flies that carry portions of the *phf7* locus, beginning at TSS1 and ending at the beginning of the open reading frame in exon 2, inserted into the same heterologous genomic location on the 3^rd^ chromosome using site specific integration. To assay for H3K9me3 accumulation on the transgene, but not at the endogenous locus, we crossed each line into a *phf7*^*Δ13*^ background. *phf7*^*Δ13*^ is a 1.81 kb intragenic deletion allele, which allows us to measure the level of H3K9me3 deposition by ChIP-qPCR at a region that is present in the transgenes but absent in *phf7*^*Δ13*^ (intronic element B, **Fig 2A**). In agreement with our previous findings, deleting the seven tandem repeats (element R) from the first intron did not interfere with ectopic H3K9me3 accumulation. (**Fig 2A and Fig 2B**). In contrast, removing elements A and R abolished H3K9me3 accumulation (**Fig 2A and Fig 2B**). These data demonstrate that cis-regulatory determinants required for H3K9me3 recruitment are within element A.

**Fig 2.**
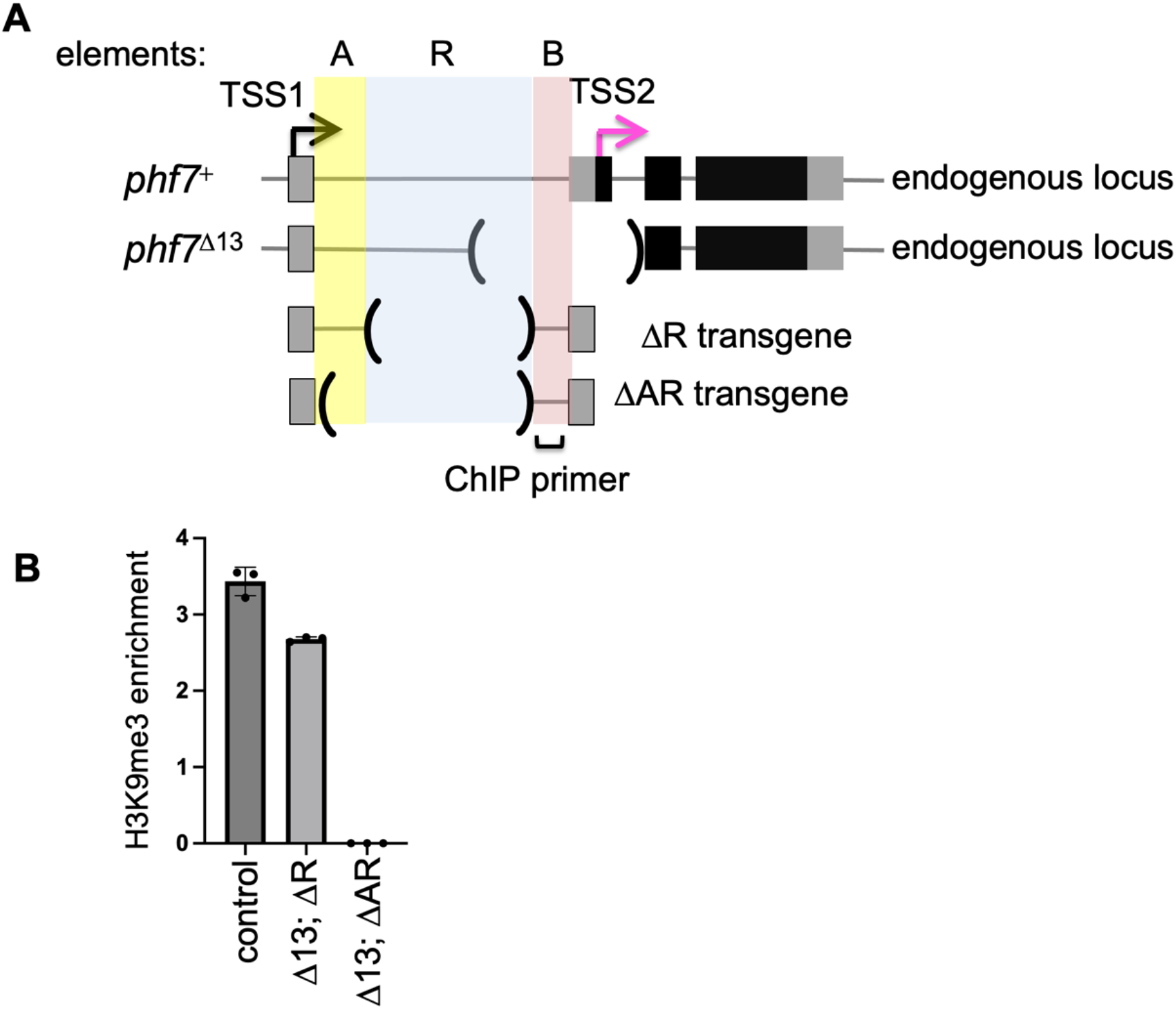
Non-coding sequences within the first intron are sufficient for H3K9me3 deposition. **(A)** Diagram of *phf7*^*+*^, *phf7*^*Δ13*^, and the genomic transgenes. **(B)** H3K9me3 accumulation on the transgenes. ChIP-qPCR measured signal, and the ChIP to input enrichment, normalized to *rp49*. Error bars represent the standard error of the mean (SD) of three biological replicates.

### Conservation of non-coding cis-regulatory elements

Cis-regulatory elements are often conserved. Thus, we might expect the essential sequences within element A to be retained in evolution. To test this prediction, we chose to examine the level of conservation between *D. melanogaster, D. simulans*, and *D. yakuba*. Although *D. simulans* and *D. yakuba* are separated from *D. melanogaster* by 5-15 million years, the *phf7* intron/exon structure is conserved (**Fig 3A**). Furthermore, RNA-seq data obtained from wild-type testis and ovaries [38,39] shows that *phf7* expression is sex specific in *D. simulans* and *D. yakuba* (**Fig 3A)**. Together these observations suggest that sex-specific transcriptional regulation through alternative TSS selection is conserved between *D. melanogaster, D. simulans*, and *D. yakuba*. When we compared the level of sequence conservation in the first intron between *D. melanogaster, D. simulans*, and *D. yakuba*, we found that only the first 235 bp of intron element A was conserved (**Fig 3B and Fig S2**). Interestingly, the first non-coding 174 bp exon also displays a high level of conservation, suggesting that additional cis-regulatory elements may be present within the first non-coding exon (**Fig 3B and Fig S2**)

**Fig 3.**
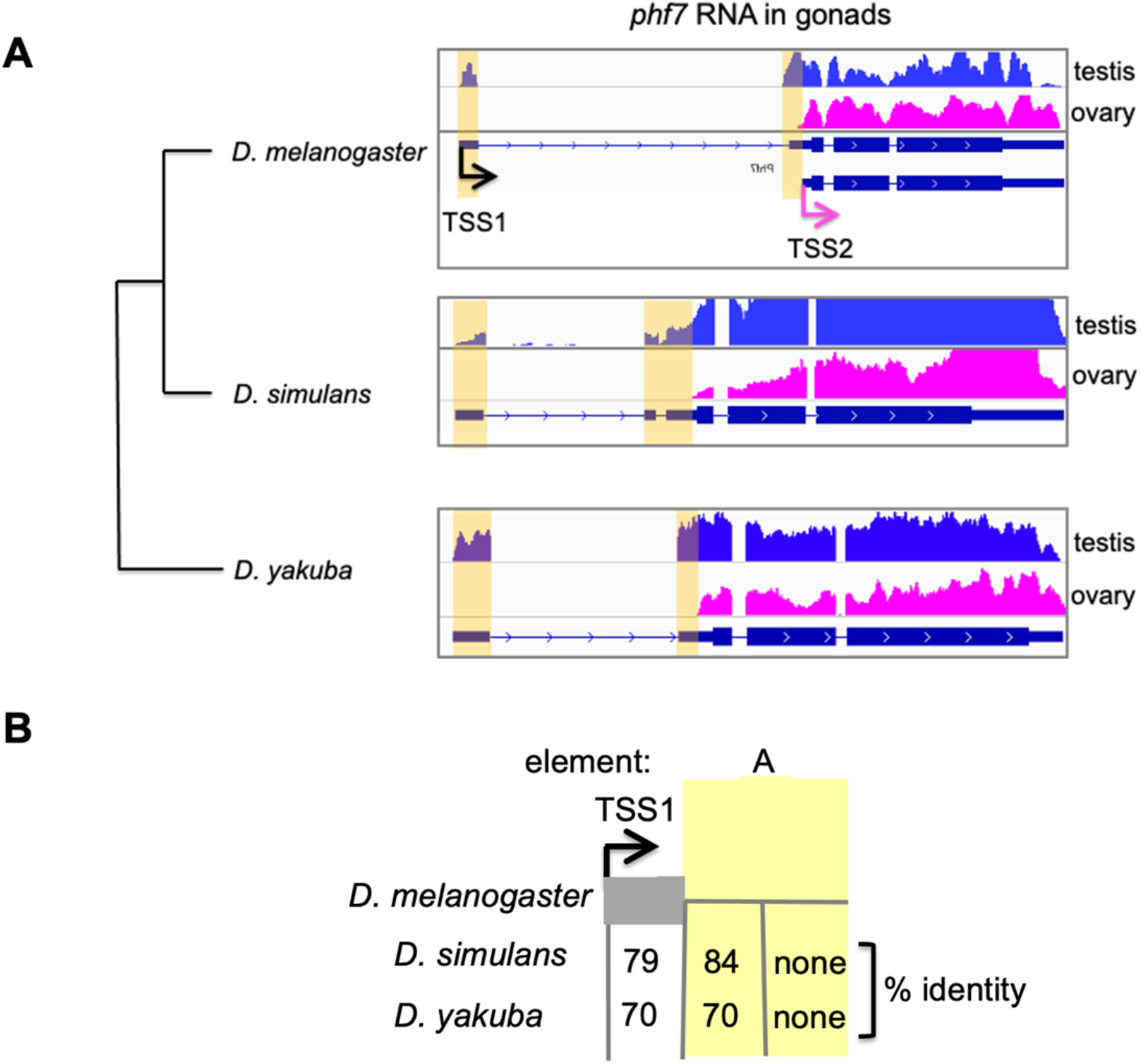
Sex specific transcriptional regulation is conserved. **(A)** Genome browser views of RNA-seq data aligned to the *phf7* locus from *D. melanogaster, D. simulans*, and *D. yakuba* ovaries and testis. **(B)** The first exon and a portion of the first intron are conserved. Pairwise alignments between the first exon and the adjacent intron were identified using the Basic Local Alignment Search Tool (BLASTn) available at (https://blast.ncbi.nlm.nih.gov). % identity to the Drosophila melanogaster *phf7* gene is shown.

### Core components of the piRNA pathway are not required for *phf7* sex specific transcriptional control

Prior studies have shown that SETDB1 is required for both TE and *phf7* silencing in germ cells [9,40–42]. Although there are no recognizable TE sequences at *phf7*, the piRNA pathway may nevertheless play a role in repressing *phf7*. To test this possibility, we analyzed published RNA-seq data sets from ovaries carrying mutations in genes encoding the dedicated piRNA pathway components, *piwi, aubergine (aub), rhino, panoramix (panx*, also known as *silenceo)*, and *nuclear RNA export factor 2 (nxf2)* [43–45]. Loss of each component causes the piRNA pathway to collapse. *piwi, aub* and *rhino* are essential for piRNA biogenesis [42,43,46]. *panx* and *nxf2*, while dispensable for piRNA biogenesis, are necessary for SETDB1 to deposit H3K9me3 marks onto TEs [45,47–51]. In agreement with our prior studies [9], only the shorter *phf7-RA* transcript is detectable in wild-type ovaries, but the longer testis specific *phf7-RC* transcript is present in *setdb1* mutant ovaries **(Fig 4)**. In contrast to the *setdb1* mutant ovaries, *piwi, aub, rhino, panx*, and *nxf2* mutant ovaries only express the shorter *phf7-RA* transcript, indicating that transcriptional regulation is not disrupted. Based on these findings we conclude that the mechanisms controlling H3K9me3 deposition onto *phf7* and TEs are distinct.

**Fig 4.**
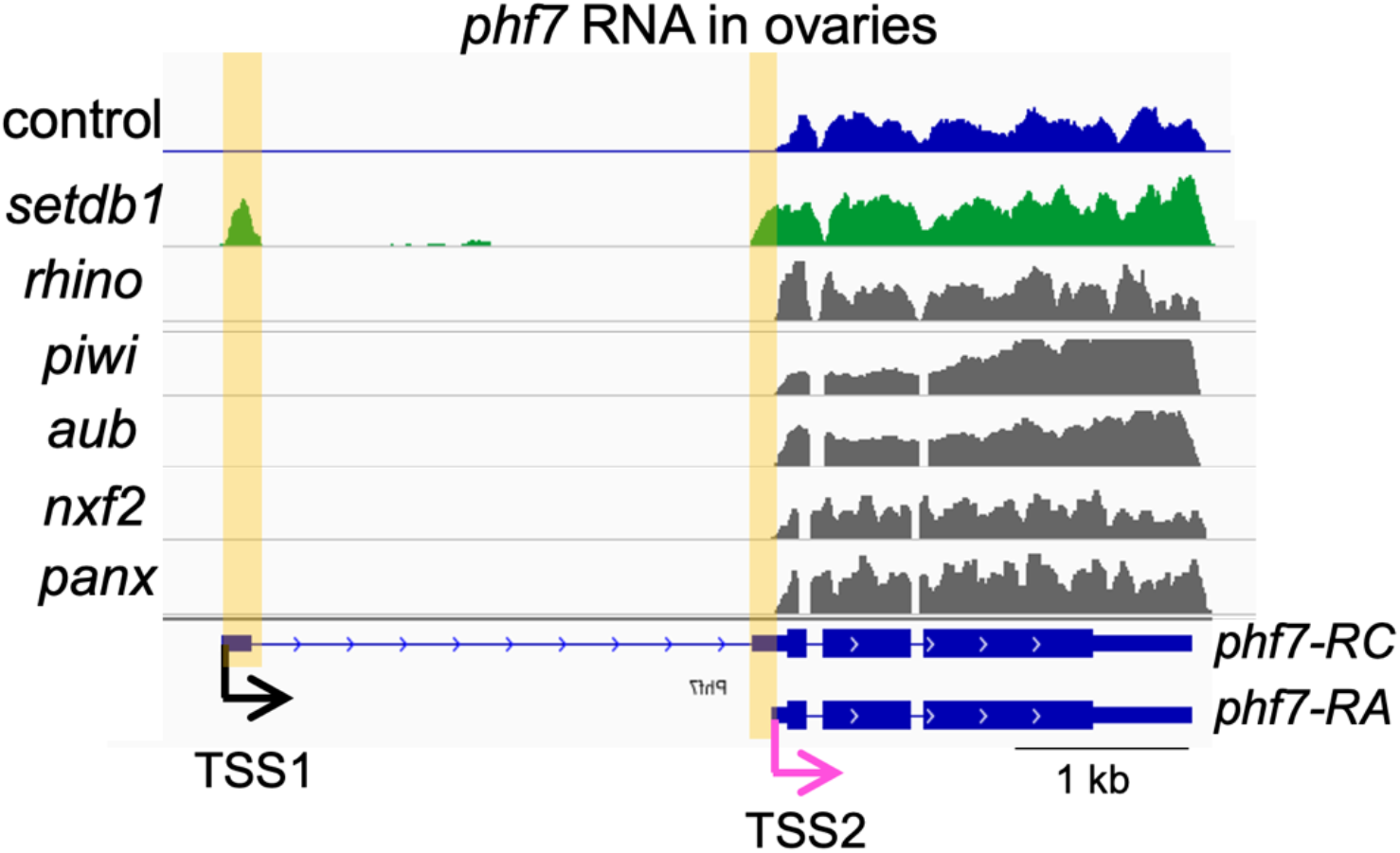
Depletion of *setdb1*, but not the dedicated piRNA-pathway components, leads to female-to-male reprogramming at the *phf7* locus. Genome browser views of the *phf7* locus. Tracks show RNA-seq reads from control and mutant ovaries aligned to the Drosophila genome (UCSC dm6). The screenshot is reversed so that the 5’ end of the gene is on the left. The RNA-seq reads that are unique to *setdb1* mutant ovaries are highlighted. In wild-type control ovaries, only the *phf7-*RA transcript is detected (TSS2, pink arrow), whereas loss of *setdb1* leads to ectopic expression of the testis specific *phf7-RC* isoform (TSS1, black arrow). In contrast, mutations in the *rhino, piwi, aub, nxf2* and *panx* genes do not disrupt sex specific *phf7* transcription. The complete mutant genotypes and accession numbers of the RNA-seq data sets are listed in Table S3.

### The ZAD-ZNF protein IDC is required for *phf7* sex specific transcriptional control

In mammalian cells, SETDB1 can be recruited to its targets by members of the large KRAB-zinc finger family of sequence specific DNA binding proteins [28–30]. The KRAB family is confined to mammals. It has been hypothesized that members of the insect specific ZAD-ZNF family might be functional analogues [52–54]. We recently tested the function of 68 of the 93 ZAD-ZNF encoding genes in female germ cells by performing an RNAi screen [55]. These data revealed that loss of CG4936 in germ cells phenocopies the ovarian defects caused by ectopic PHF7 expression. CG4936, which we name Identity crisis (IDC), is a 521 amino acid (aa) protein that contains an N-terminal ZAD (zinc finger associated domain, 22-95 aa), an unstructured linker region, and a C-terminal domain that includes an array of 5 C2H2 zinc fingers (386-491 aa). Studies of other ZAD-ZNF proteins suggest that the ZAD mediates protein-protein interactions and the C2H2 zinc fingers bind DNA [56–60]. We, therefore, hypothesize that IDC is required for *phf7* repression.

To investigate the possibility that loss of *idc* in female germ cells disrupts *phf7* repression, we first used RT-qPCR to assay for the presence of the testis specific *phf7-RC* transcript. Germ cell specific knock down (GLKD) was achieved using an inducible RNA interference (RNAi) transgene and the germ cell specific *nos-GAL4* driver. To rule out indirect effects arising from RNAi knockdown, we also knocked down *white*, which is not expressed in germ cells, as a negative control. Using primer pairs capable of detecting *phf7-RC*, we found that in contrast to control ovaries, *phf7-RC* is detectable in *idc* GLKD mutant ovaries (**Fig 5A**). We then asked whether ectopic *phf7-RC* expression correlated with ectopic PHF7 protein expression. Previous work has shown that a BAC transgene that encodes an N-terminally HA-tagged PHF7 protein (HA-PHF7) is a faithful reporter of the sex specific protein expression pattern of PHF7 [37,61]. We found that, in contrast to wild-type ovaries, the HA-PHF7 protein is detectable in the cytoplasm and the nucleus of the *idc GLKD* mutant ovaries (**Fig 5B and Fig 5C**). We conclude that IDC is required for *phf7* regulation.

**Fig 5.**
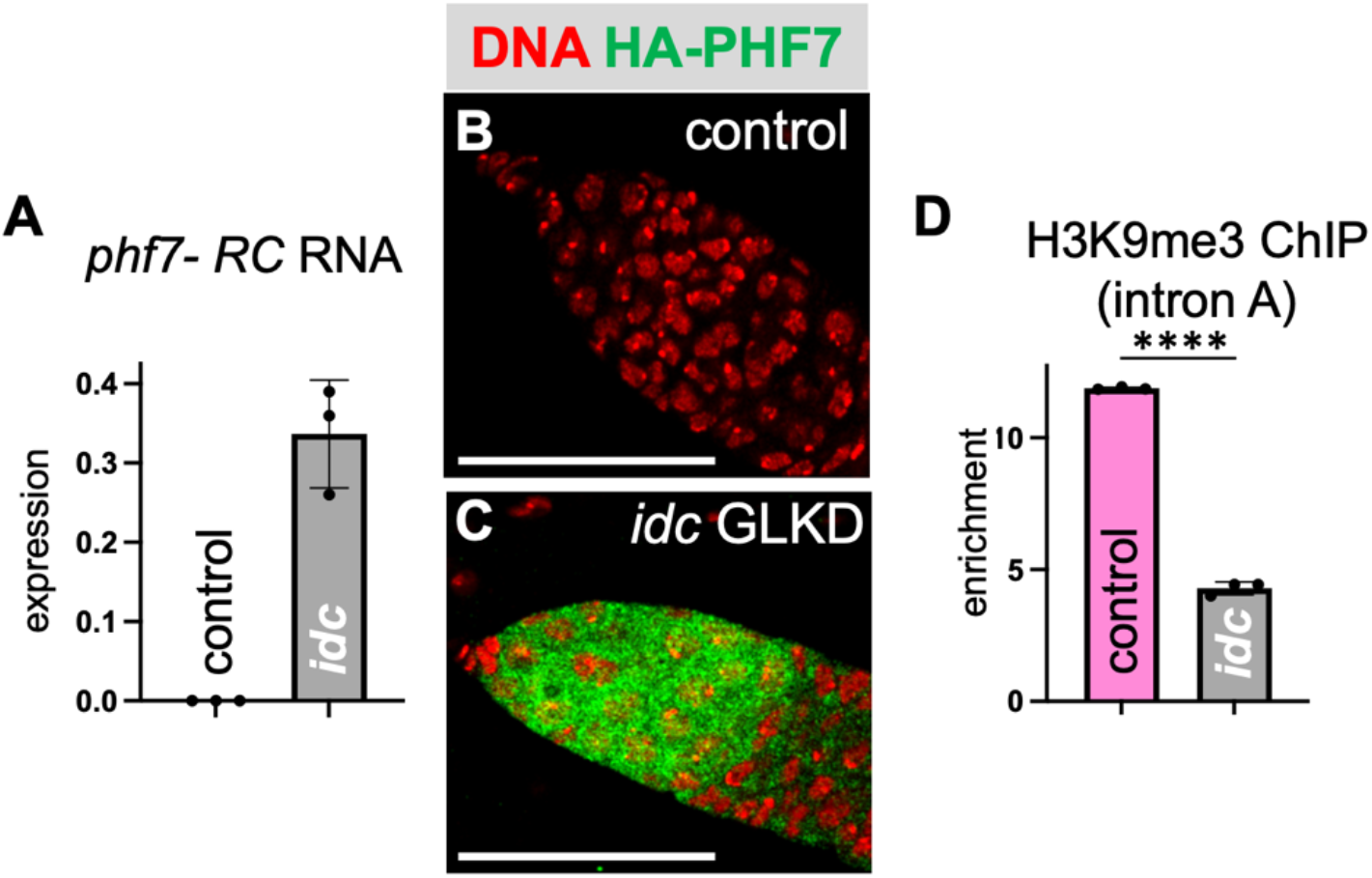
*idc* germ cell specific knock-down disrupts *phf7* regulation. **(A)** *idc* GLKD germ cells (*nos>idc-RNAi*) express the testis *phf7-RC* transcript. RT-qPCR measures expression normalized to the total level of *phf7* in RNA extracted from control *(nos>white-RNAi*) and *idc* GLKD (*nos>idc-RNAi*) ovaries. The histograms show the mean ± SD of three biological replicates. **(B, C)** *idc* GLKD germ cells inappropriately express the PHF7 protein. Confocal images of ovaries were dissected from control (*nos>white-RNAi*) and mutant (*nos>idc-RNAi*) females carrying an HA-PHF7 transgene and stained for HA (green) and DNA (red). Scale bar 50μm. **(D)** H3K9me3 accumulation at the endogenous *phf7* locus is reduced in *idc* GLKD ovaries. ChIP-qPCR measured H3K9me3 signal at the *phf7* locus (primer in region A) in control (*nos>white-RNAi)* and mutant (*nos>idc-RNAi*) ovaries. ChIP to input enrichment is normalized to *rp49*. The histogram shows the mean ± SD of three biological replicates. A two-tailed Student’s t-test estimates statistical significance where \*\*\*\**p*<0.0001.

To test whether *idc* plays a role in controlling H3K9me3 deposition, we compared the amount of H3K9me3 accumulation within element A in wild-type and mutant ovaries using ChIP-qPCR. We found that H3K9me3 was significantly reduced in *idc* GLKD ovaries compared to control ovaries (*p*<0.0001; **Fig 5D**). Based on these studies, we conclude that IDC regulates *phf7* transcription by promoting H3K9me3 deposition.

### IDC is a nuclear protein that associates with DNA in ovaries

To understand the role of IDC in H3K9me3 deposition, we first determined its localization pattern in the adult ovary. We used a genomic fosmid transgene for these studies that encodes a C-terminally GFP-tagged IDC protein (IDC-GFP). We inferred that the GFP tag does not interfere with *idc* function because the *idc-GFP* rescues the *idc*^*1*^ lethal phenotype (see materials and methods). Each ovary comprises 16-20 ovarioles that contain germ cells spanning the range of maturity from germline stem cells (GSCs) at the anterior end to fully mature eggs at the posterior end [62]. We found that IDC-GFP is detectable in the nucleus of several different cell types within the egg chambers, including nurse cells, oocytes, and somatic follicle cells (**Fig 6A**). IDC-GFP is tightly associated with the polytene chromosomes in the nurse cells, consistent with its presumed DNA binding activity (**Fig 6B**).

**Fig 6.**
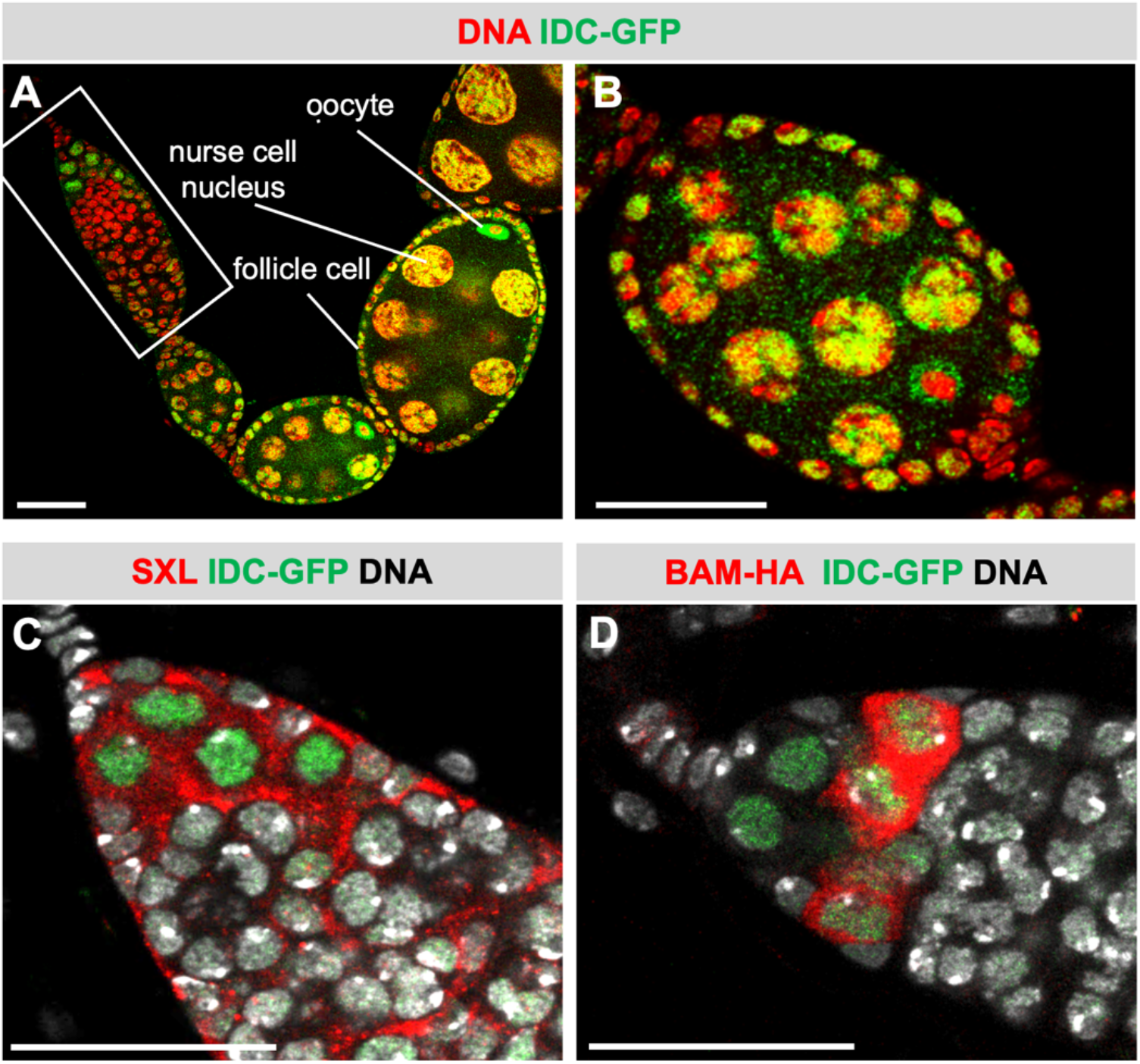
IDC is nuclear protein expressed in both somatic and germline cells. **(A)** Confocal image of an ovariole dissected from a female carrying the IDC-GFP rescue transgene stained for GFP (green) and DNA (red). The germarium at the anterior end of the ovariole is boxed in. An oocyte, a nurse cell, and a somatic follicle cell are indicated. Scale bar 25μm. **(B)** Confocal image of an early-stage egg chamber dissected from a female carrying the IDC-GFP transgene stained for GFP (green) and DNA (red). Scale bar 25μm. **(C)** Confocal image of the anterior end of a germarium dissected from a female expressing the IDC-GFP fusion protein and co-stained for GFP (green), DNA (white), and SXL (red). Scale bar 25 μm. **(D)** Confocal image of the anterior end of a germarium dissected from a female expressing the IDC-GFP and the BAM-HA fusion proteins co-stained for GFP (green), DNA (white), and HA (red). Scale bar 25μm.

In the germarium, at the anterior end of the ovariole, we found that the IDC-GFP protein accumulates to high levels in the nucleus of only 3 to 6 germ cells (boxed region; **Fig 6A)**. This expression pattern is like that described for the female specific sex determination protein SEX-LETHAL (SXL), which is also required for *phf7* H3K9me3 silencing [9,63,64]. Indeed, co-staining experiments reveal that SXL accumulates in the cytoplasm of all IDC-GFP expressing germ cells (**Fig 6C**). Studies have shown that SXL accumulates in GSCs and their immediate daughter cells. When GSCs divide, the daughter cells at the tip remain a GSC, while the more posterior daughter cells, called a cystoblast (CB), express the BAG OF MARBLES protein (BAM). In agreement with published studies, BAM expression, assessed with a transgene that encodes a C-terminally HA-tagged BAM protein expressed from a *bam* promoter is readily detectable in the cytoplasm of just a few germ cells [65]. Co-staining experiments reveal that IDC-GFP is expressed in the GSCs where BAM-HA is not expressed as well as in the adjacent two to three cells where BAM-HA is detectable (**Fig 6D**). Based on these observations, we conclude that the nuclear IDC protein is expressed in the GSCs and their immediate daughter cells.

### IDC binds to sequences within the male specific exon

IDC contains five zinc fingers suggesting that it has sequence specific DNA binding activity. We speculated that IDC might promote H3K9me3 deposition by binding to the *phf7* locus. To test this prediction by ChIP-qPCR, we designed primers along the length of the conserved noncoding sequences in the first exon and the intron. These experiments revealed that the IDC-GFP tagged protein associates with chromatin within the first non-coding exon, but not outside of it (**Fig 7A**). Collectively, our studies indicate that IDC regulates *phf7* directly by serving as an H3K9me3 guidance factor.

**Fig 7.**
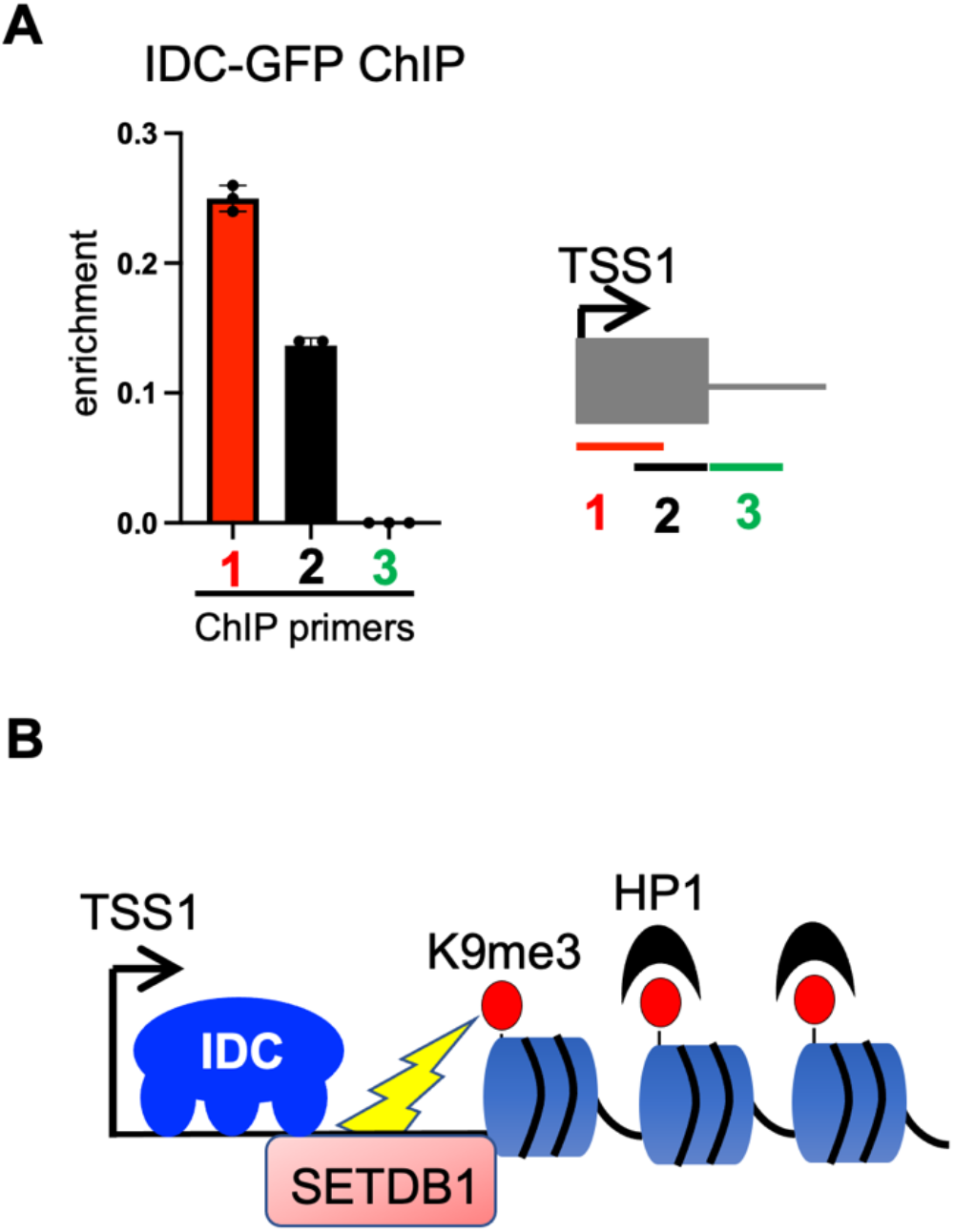
IDC binds to *phf7*. (A) ChIP-qPCR shows that IDC binds to the *phf7* first exon. Left: The histogram shows the mean ± SD of three biological replicates. Right: Cartoon showing the location of the primers used for ChIP-qPCR. **(B)** Model of H3K9me3-mediated silencing of *phf7*. IDC binds to *phf7*, directing H3K9me3 deposition by SETDB1 H3K9 methytransferase. HP1 binds to H3K9me3, resulting in transcriptional silencing.

## Discussion

SETDB1-controlled H3K9 methylation plays an essential role in securing female germ cell fate by silencing lineage-inappropriate *phf7* transcription [9,36]. SETDB1 is also required for TE silencing, where a piRNA-guided mechanism guides H3K9me3 deposition [40–42]. In this work, we establish that *phf7* is silenced by a piRNA-independent mechanism and discover that regulation depends on IDC, a previously uncharacterized member of the ZAD-ZNF family of DNA binding proteins. Regulation appears direct, as IDC is required for H3K9me3 deposition and localizes to a conserved region within the *phf7* locus. Collectively, our data establish that the sequence specific DNA binding protein IDC guides the installation of the H3K9me3 mini-domain onto the *phf7* locus, thereby preventing accidental female-to-male reprogramming. In addition to extending our understanding of how female germ cell fate is maintained, our studies provide one of the first examples of a ZAD-ZNF protein guiding H3K9me3-mediated gene silencing.

Our work is consistent with a simple model in which IDC recruits the SETDB1 H3K9me3 methyltransferase to *phf7* (**Fig 7B**). However, we could not detect an interaction between IDC and SETDB1 in co-immunoprecipitation experiments. It is, therefore, possible that another as yet undiscovered protein recruits SETDB1 to *phf7* and that IDC acts at a later step in the pathway to ensure the sequence specific establishment of the H3K9me3 mini domain.

The IDC protein is also expressed in male germ cells, suggesting that an accessory protein is required to confer female specificity to *phf7* silencing. One attractive contender is *stonewall (stwl)*. STWL is a female specific heterochromatin-associated protein that acts as a transcriptional repressor *in vitro* [66], associates with SETDB1 in yeast [67] and localizes to the *phf7* locus in S2 cells [68]. Importantly, loss of *stwl* in ovaries leads to the inappropriate expression of the testis *phf7* transcript [68]. Together, these studies raise the possibility that *stwl* collaborates with IDC to silence *phf7* in female germ cells.

We have previously established that the H3K9me3 reader protein, HP1a, is essential for *phf7* silencing [9]. A requirement for HP1a is not surprising, as HP1a drives chromatin compaction and transcriptional silencing [69]. HP1a also recruits H3K9 methyltransferases, enabling the spreading of the H3K9me3 repressive domain through a positive feedback mechanism. The spread of H3K9me3 over the three kb region within the *phf7* gene likely occurs by an analogous mechanism. At the *phf7* locus, however, the H3K9me3 domain does not extend into the open reading frame or the neighboring genes. Therefore, more work is needed to understand what stops the spreading of this repressive chromatin domain.

The SETDB1-controlled H3K9 methylation silencing pathway in female germ cells is not restricted to *phf7*. Genome-wide H3K9me3 profiling in wild-type and mutant ovaries has shown that SETDB1 silences two classes of protein-coding genes. One type includes genes usually expressed in the testis [9]. A second class includes genes that are typically expressed in undifferentiated female germ cells but are silenced once the oocyte is specified, such as *ribosomal protein S19b* [70]. These few examples of context dependent H3K9me3 gene silencing suggest a finely tuned guidance mechanism in the female germline. It will be interesting to explore whether other members of the ZAD-ZNF gene family serve as H3K9me3 guidance factors.

In summary, by focusing on a single biologically relevant gene, we discovered a putative DNA binding protein that guides the installation of a H3K9me3 repressive domain onto a protein-coding gene. These findings are reminiscent of how TEs can be silenced in mammals wherein members of the KRAB-ZNF protein family recruit SETDB1 to establish epigenetic repression [28–31]. Interestingly, the KRAB-ZNFs are vertebrate specific, and the ZAD-ZNFs are insect specific. Yet, both gene families exhibit similar patterns of species-specific gene expansion and diversification. These observations have fueled the speculation that ZAD-ZNFs and KRAB-ZNFs perform similar functions [52–54]. In fact, IDC’s closest human relative is ZNF75D, a KRAB-ZNF protein of unknown function. Whether this or other members of the KRAB-ZNF protein family are required for tissue specific H3K9me3 gene silencing will be an exciting area of further exploration.

## Materials and Methods

### *Drosophila* stocks and culture conditions

Fly strains were kept on standard media at 25 °C unless otherwise noted. Wild-type control flies were from the lab *y*^*1*^ *w*^*1*^ stock. Stocks used to report on protein gene activity include the HA-tagged *phf7* transgene, *PBac{3XHA-PHF7}* [37]; the GFP-tagged *idc* transgene, *FlyFos{fTRG00376*.*sfGFP-TVPTBF}* (VDRC #318556) described in [71] and the HA-tagged *bam* transgene, *P{Bam::HA}* [65]. *phf7*^*Δ13*^, *phf7*^*ΔR*^, the *idc*^*1*^ null allele, and the ΔR and ΔAR chromatin reporters were generated for this study, as described below.

Crosses for knock-down studies were set up at 29°C, and adults were aged 3-5 days before gonad dissection. Knock-down studies were carried out with the following germline-optimized lines generated by the *Drosophila* Transgenic RNAi Project [72]: *idc*-*P{TRiP*.*HMC05569}* (BDSC #64550), and the control *w-P{TRiP*.*GL00094}* (BDSC #35573). Expression of the UAS-RNAi transgenic lines was driven by the third chromosome *P{GAL4::VP16-nos*.*UTR}* insertion (BDSC #4937), which is strongly expressed in germ cells.

### Generation of the *phf7*^*ΔR*^ allele

We generated the *phf7*^*ΔR*^ allele in two steps. First, CRISPR was used to replace the repeats within the first intron with a 3XP3::DsRed cassette by Rainbow Transgenic Flies, Inc. The deletion was generated with the following guide RNAs: agttaaaaaaaatcaatcgatgg and cgcagcgattgaatgttaatggg. 1 kb homology arms were generated through PCR and cloned into the pHD-dsRed-attP (Addgene #51019) [73]. Guide RNAs and the donor vector were co-injected into *nos-Cas9* embryos by Rainbow Transgenic Flies. Flies were screened for DsRed expression in the eyes, and sequence verified for accuracy. *phf7*^*ΔR*^ was generated by removing the DsRed cassette. Homozygous *phf7*^*ΔR+dsRED*^ females were crossed to a Cre-expressing fly line (BDSC #1092), and the male progeny screened for the loss of dsRED expression in the eyes, followed by sequence confirmation of precise tag excision.

### Generation of the *phf7*^*Δ13*^ allele

The *phf7*^*Δ13*^ allele was generated by P-element mediated imprecise excision as follows: females homozygous for the P-element insertion P{EPgy2}Phf7^EY03023^ (BDSC #15894) were crossed with the Δ2-3 transposase expressing line (BDSC #3664). Potential excision lines were established from the male progeny of the F1 females that had lost the *white*^*+*^ marker carried by the P element. Stocks that carried deletions were identified by PCR. The exact breakpoints were determined by DNA sequencing the PCR amplified DNA fragments.

### Generation of the ΔR and ΔAR chromatin transgenic reporter lines

Constructs were generated by VectorBuilder’s custom cloning services (https://en.vectorbuilder.com). The transgenes were constructed in their “user-defined promoter” modification of the pUASTattB expression vector. The transgenic constructs were sent to Rainbow Transgenic Flies Inc. for *phi-C31* catalyzed integration into the 65B2 PBac{y[+]-attP-3B}VK00033 site. Transgenic flies were sequence verified for accuracy.

### Generation of the *idc*^*1*^ null allele and rescue by the GFP-tagged *idc* transgene

The *idc*^*1*^ allele was generated by *in vivo* CRISPR mutagenesis [74]. Females expressing two sgRNAs under control of the GAL4/UAS system, P{HD_CFD00598} (VDRC #341525), were crossed at 25°C to M{vas-Cas9}; P{GAL4::VP16-nos.UTR} (BDSC #55821 + #4937) males. The *vas-Cas9; nos>gRNA* male offspring were crossed to a third chromosome balancer line to isolate and balance each putative *idc* allele in the next generation. *idc* alleles were identified by the failure to complement Df(3R)ED6027 (BDSC #9479) and checked for the presence of the predicted indel by PCR.

We found that *idc*^*1*^*/Df(3R)ED6027* animals do not survive to adulthood (n>100). To verify that the GFP-tagged *idc* transgene, PBac{fTRG00376.sfGFP-TVPTBF} (VDRC #318556), rescues *idc*^*1*^, we first made the double mutant *idc*^*1*^, *idc-GFP*. To test for rescue, we crossed *idc*^*1*^, *idc-GFP/TM3* to *Df(3R)ED6027/TM3* males. This cross yielded 84% (31/37) of the expected *idc*^*1*^, *idc-GFP/ Df(3R)ED6027* progeny, all of which were fertile.

### qRT-PCR and data analysis

RNA was extracted from dissected gonads using TRIzol (Thermo Fisher, cat# 15596026). Quantity and quality were measured using a NanoDrop ND-1000 spectrophotometer. The RNA was treated with DNase RQ1 (Promega, cat# M6101). cDNA was generated by reverse transcription using a SuperScript First-Strand Synthesis System for RT-PCR kit (Thermo Fisher, cat# 11904018) using random hexamers. qPCR was performed using *Power* SYBR Green PCR Master Mix (Thermo Fisher, cat# 4368706) with an Applied Biosystems QuantStudio 3 Real-Time PCR system. PCR steps were as follows: 95°C for 10 minutes followed by 40 cycles of 95°C for 15 seconds and 60°C for 1 minute. Melt curves were generated with the following parameters: 95°C for 15 seconds, 60°C for 1 minute, 95°C for 15 seconds, and 60°C for 15 seconds. Measurements were taken in biological triplicate with two technical replicates each. Relative transcript levels were calculated using the 2-ΔΔCt method [75]. Using GraphPad Prism software, P values were calculated using unpaired two-tailed Student’s t-tests. The primers used for measuring RNA levels are listed in **Table S1**.

### ChIP-qPCR and data analysis

ChIP was performed as described in [9] with some modifications. Buffers are listed in **Table S2**. Briefly, ovaries from 200 1-2 day old adults were dissected in PBSP, homogenized with a pellet pestle, and crosslinked with 1.8% methanol-free formaldehyde (Thermo Fisher, cat# 28906) for 5 minutes. Fixation was quenched for 5 minutes by adding glycine to a final concentration of 225mM. The solution was removed, and the pellet was washed in PBSP 3 times. Samples were resuspended in 500μl PBSP and stored at -80°C. Samples were thawed, centrifuged, and resuspended in 1ml lysis buffer 1. Samples were homogenized using sterile homogenizing beads in a Bullet Blender Homogenizer at 4°C for six cycles of 30 seconds on and 1 minute off. Following centrifugation, cell pellets were washed at 4°C for 10 minutes in lysis buffer 1, centrifuged, washed for 10 min at 4°C in lysis buffer 2, centrifuged and then resuspended in 750μl lysis buffer 3. Chromatin was sheared to 100–500 base pairs at 4°C with a Diagenode BioRuptor Pico for 15 cycles of 30 seconds on, and 30 seconds off.

For H3K9me3 ChIP, the chromatin lysates were precleared with a 1:1 mix of protein A/G Dynabead magnetic beads (Thermo Fisher, cat# 10001D and 10003D) for 1 hour at 4°C and then incubated overnight at 4°C with 1:1 mix of protein A/G Dynabead magnetic beads conjugated to H3K9me3 antibody (Abcam, cat# 8898 RRID: AB_306848).

For GFP ChIP, the chromatin lysates were precleared with Chromotek Magnetic Binding Control Agarose Beads (Proteintech, cat# bmab) for 1 hour at 4°C, and then incubated overnight at 4°C with Chromotek GFP-Trap Magnetic Agarose beads (Proteintech, cat# gtma).

Following immunoprecipitation, the samples were washed six times with ChIP-RIPA buffer, two times with ChIP-RIPA/500 buffer, two times with ChIP-LiCl buffer, and twice with TE buffer. DNA was eluted and reverse crosslinked from beads in 200μl elution buffer overnight at 65°C. Following RNaseA and proteinase K treatment, DNA was recovered by phenol-chloroform extraction and used for qPCR.

qPCR was performed using *Power* SYBR Green PCR Master Mix (Thermo Fisher, cat# 4367659) with an Applied Biosystems QuantStudio 3 Real-Time PCR system. PCR steps were as follows: 95°C for 10 minutes followed by 40 cycles of 95°C for 15 seconds and 60°C for 1 minute. Melt curves were generated with the following parameters: 95°C for 15 seconds, 60°C for 1 minute, 95°C for 15 seconds, and 60°C for 15 seconds. Primers as listed in **Table S1**. ChIP experiments were performed on three independent biological samples. qPCR measurements on each sample were performed on at least two technical replicates. For H3K9me3 ChIP-qPCR, the ChIP to input enrichment (presented as percent input) is normalized to a negative control genomic region in the *rp49* locus. Statistical analysis was carried out by the GraphPad Prism software. The P values were calculated using unpaired two-tailed Student’s t tests.

### Immunofluorescence and image analysis

Females were aged 3-5 days before gonad dissection. Ovaries were fixed and stained according to standard procedures with the following primary antibodies: mouse anti-*Drosophila* SXL (1:100, Developmental Studies Hybridoma Bank, cat# M18, RRID: AB_528464), rabbit anti-GFP (1:2500, Thermo Fisher, cat# A-11122, RRID: AB_221569) or FITC conjugated goat anti-GFP (1:750, Abcam, cat# ab6662, RRID: AB_305635), and rat anti-HA high affinity (1:500, Sigma, cat# 11867423001, RRID: AB_390919). The following secondary antibodies were used at 1:200: Alexa Fluor 555 goat anti-rat (Thermo Fisher, cat# A-21434, RRID: AB_2535855), FITC goat anti-mouse (Jackson Immunoresearch Laboratories, cat# 115-095-003, RRID: AB_2338589), or FITC goat anti-rabbit (Jackson Immunoresearch Laboratories, cat# 111-095-003, RRID: AB_2337972). TO-PRO-3 Iodide carbocyanine monomer nucleic acid stain (1:1000, Thermo Fisher, cat# T3605) was used to stain DNA. Images were taken on a Leica SP8 confocal with 1024×1024 pixel dimensions, a scan speed of 600 Hz, and a frame average of 3. Sequential scanning was done for each channel, and three Z-stacks were combined for each image. Processed images were compiled with Microsoft PowerPoint.

### Analysis of publicly available RNA-seq data sets

**Table S3** lists the sources of the publicly available RNA-seq data generated from wild-type and mutant *Drosophila* tissues. The data were downloaded from the SRA database at NCBI and analyzed using the tools available on the Galaxy web platform (https://usegalaxy.org/). FastQC assessed read quality and STAR was used to align the reads to the *Drosophila* reference genome (UCSC dm6). Screenshots of the expression data displayed on The Integrated Genome Viewer (IGV) are shown.

*D. simulans* and *D. yakuba phf7* orthologs were identified using the ortholog database at www.flybase.org. RNA-seq data from wild-type *D. simulans* and *D. yakuba* gonads was downloaded from the SRA database at NCBI (**Table S4**). We uploaded the sequencing data to the Galaxy web platform (https://usegalaxy.org/), assessed read quality with FastQC and used STAR to align the reads to their respective reference genomes (listed in **Table S4**). Browser screenshots of the expression data are displayed on the Integrated Genome Viewer (IGV).

### DNA sequence alignments

Multiple sequence alignments were performed in SnapGene (https://www.snapgene.com) using MUSCLE with default options. Pairwise sequence alignments were performed in BLASTn (https://blast.ncbi.nlm.nih.gov).

## Acknowledgments

We would like to thank the members of the Cleveland fly community for helpful discussions, Jane Heatwole for fly food, and FlyBase as an essential resource. Stocks obtained from the Bloomington (NIH P40D018537) and The Vienna *Drosophila* Stock Centers were used in this study. Imaging was performed using equipment purchased through NIH Grant S10-OD016164, housed in the CWRU School of Medicine Imaging Core.

## Author contributions

HS conceived this study. LSK and HS generated reagents, designed and executed the experiments, and interpreted the data. MS performed the evolutionary analysis. HS wrote the manuscript with input from LSK and MS.

## Declaration of interests

The authors declare no competing interests

## Availability of data and material

All genomic sequence and expression data were obtained from public databases and are described in **Table S3** and **S4**. *Drosophila* strains are available upon request.

## Figures

**Figure S1.**
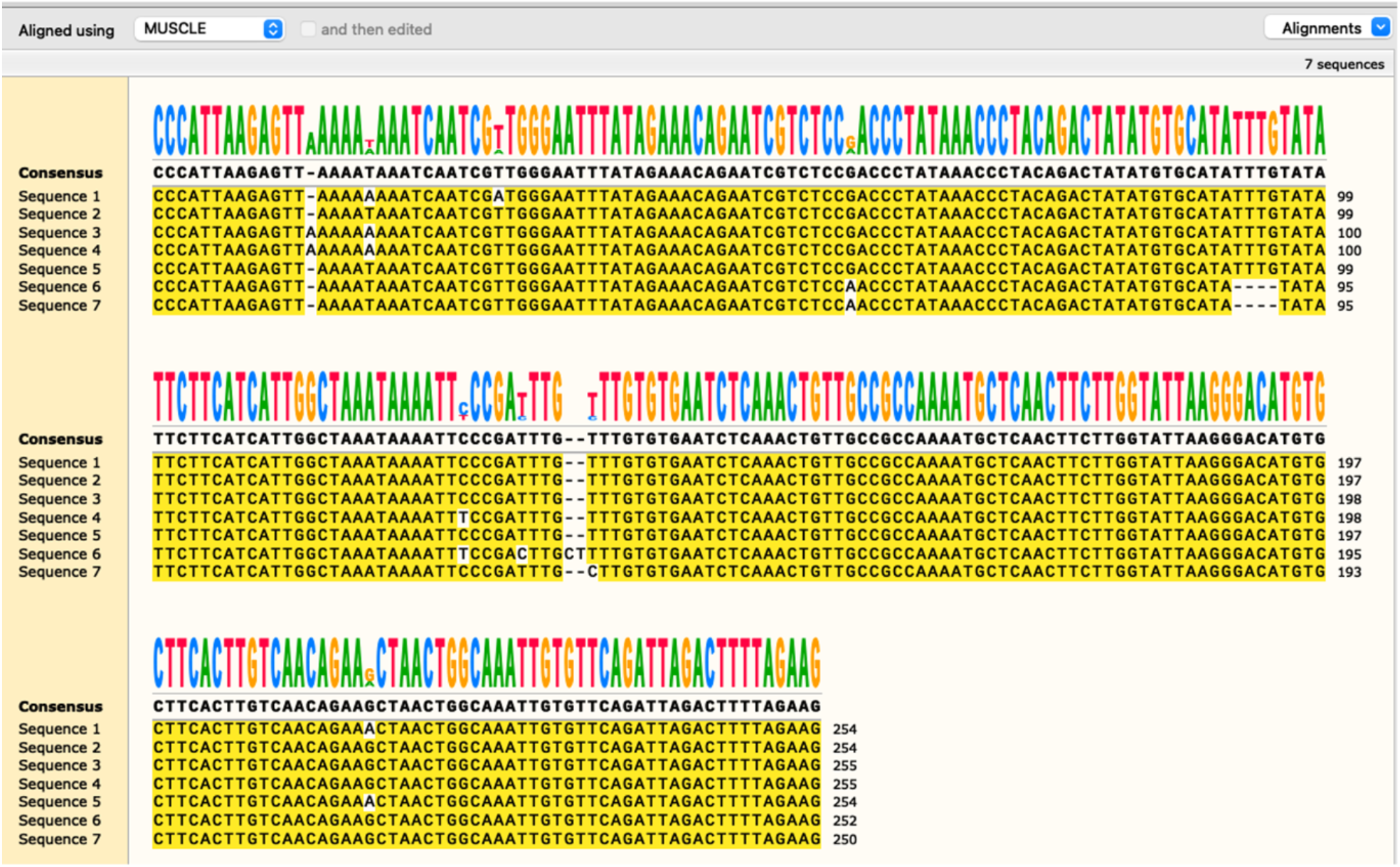
The first intron of *phf7* contains seven copies of an ∼250 bp DNA sequence. Sequence alignment of the 7 ∼250 bp sequences generated with the multiple sequence alignment tool MUSCLE embedded in the SnapGene software (snapgene.com).

**Figure S2.**
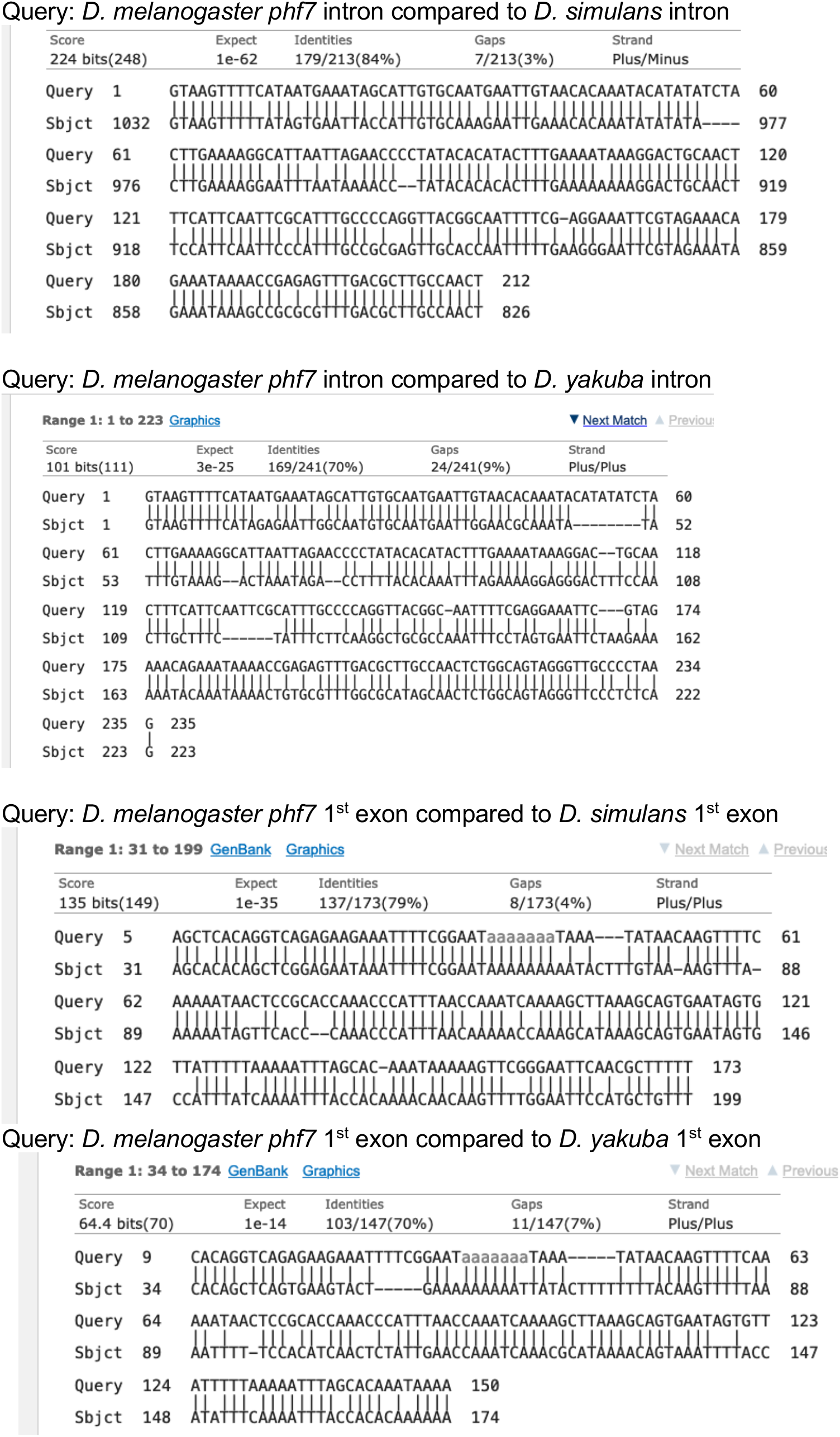
The first exon and a portion of the first intron are conserved between *D. melanogaster, D. simulans* and *D. yakuba*. Pairwise alignments between the first exon and the adjacent intron were identified using the Basic Local Alignment Search Tool (BLASTn) available at (https://blast.ncbi.nlm.nih.gov).

**Table S1.**
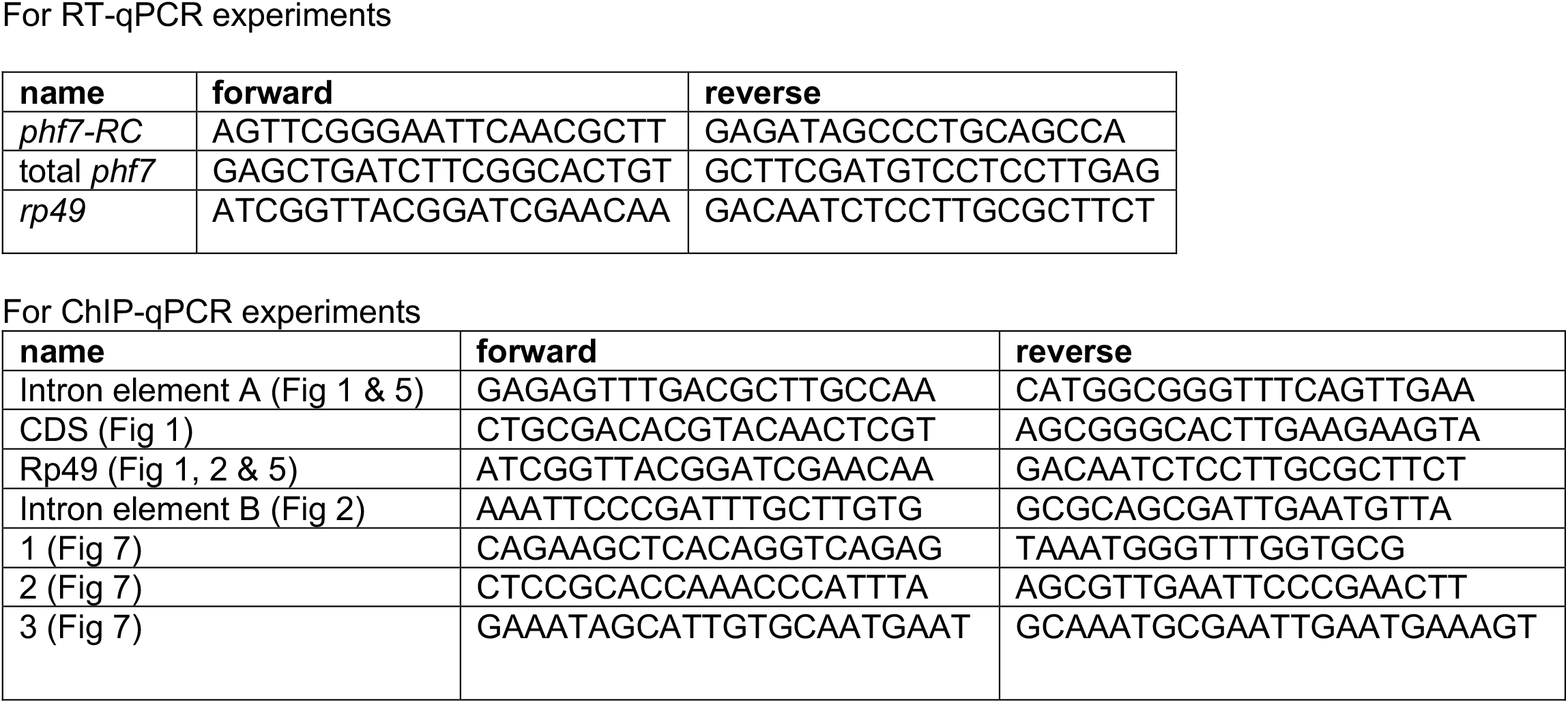
Primers.

**Table S2.**
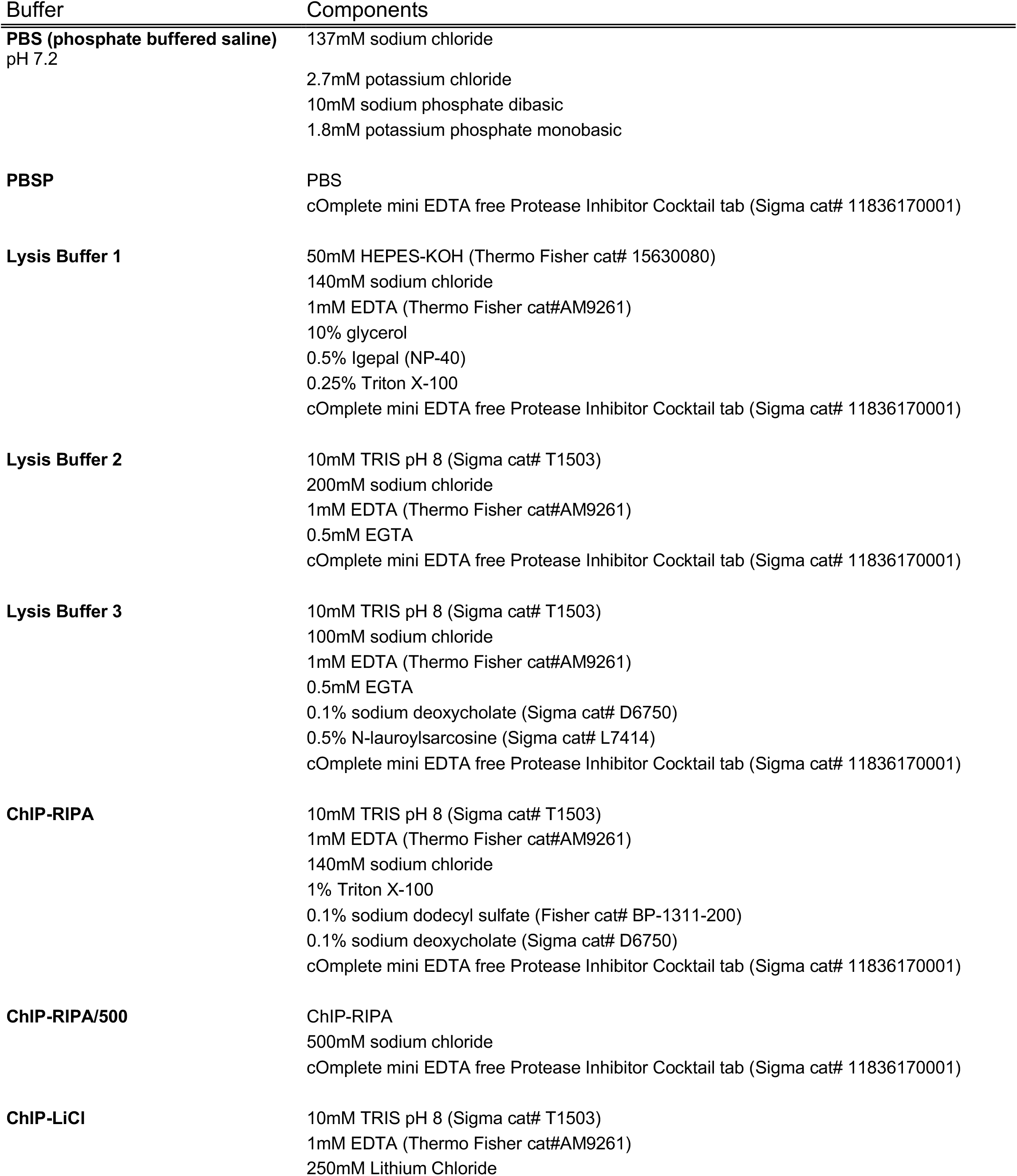
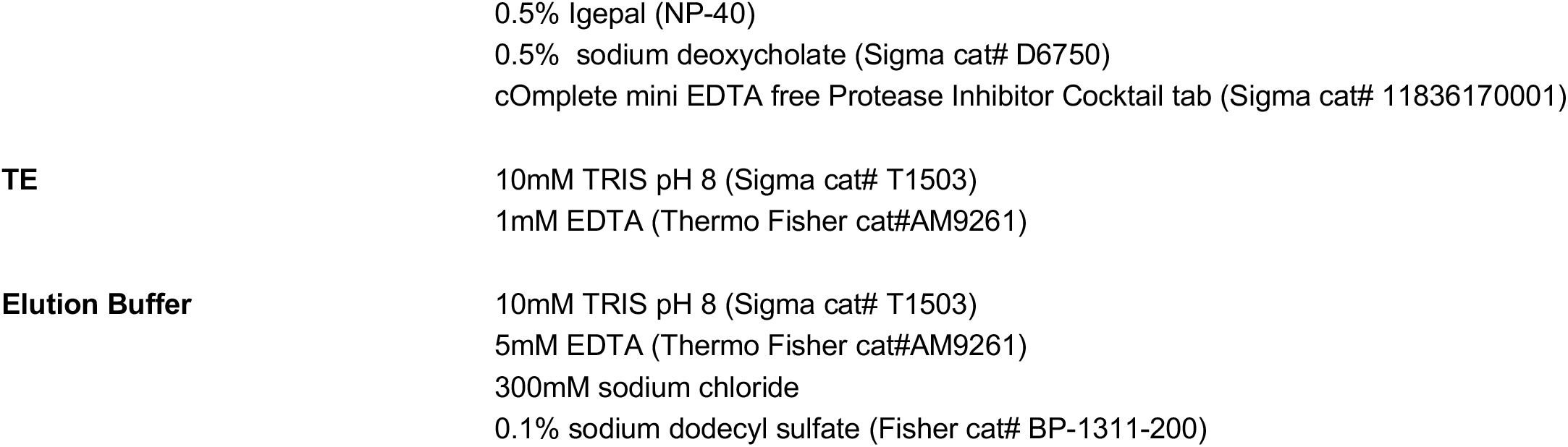
Buffers for ChIP.

**Table S3.**
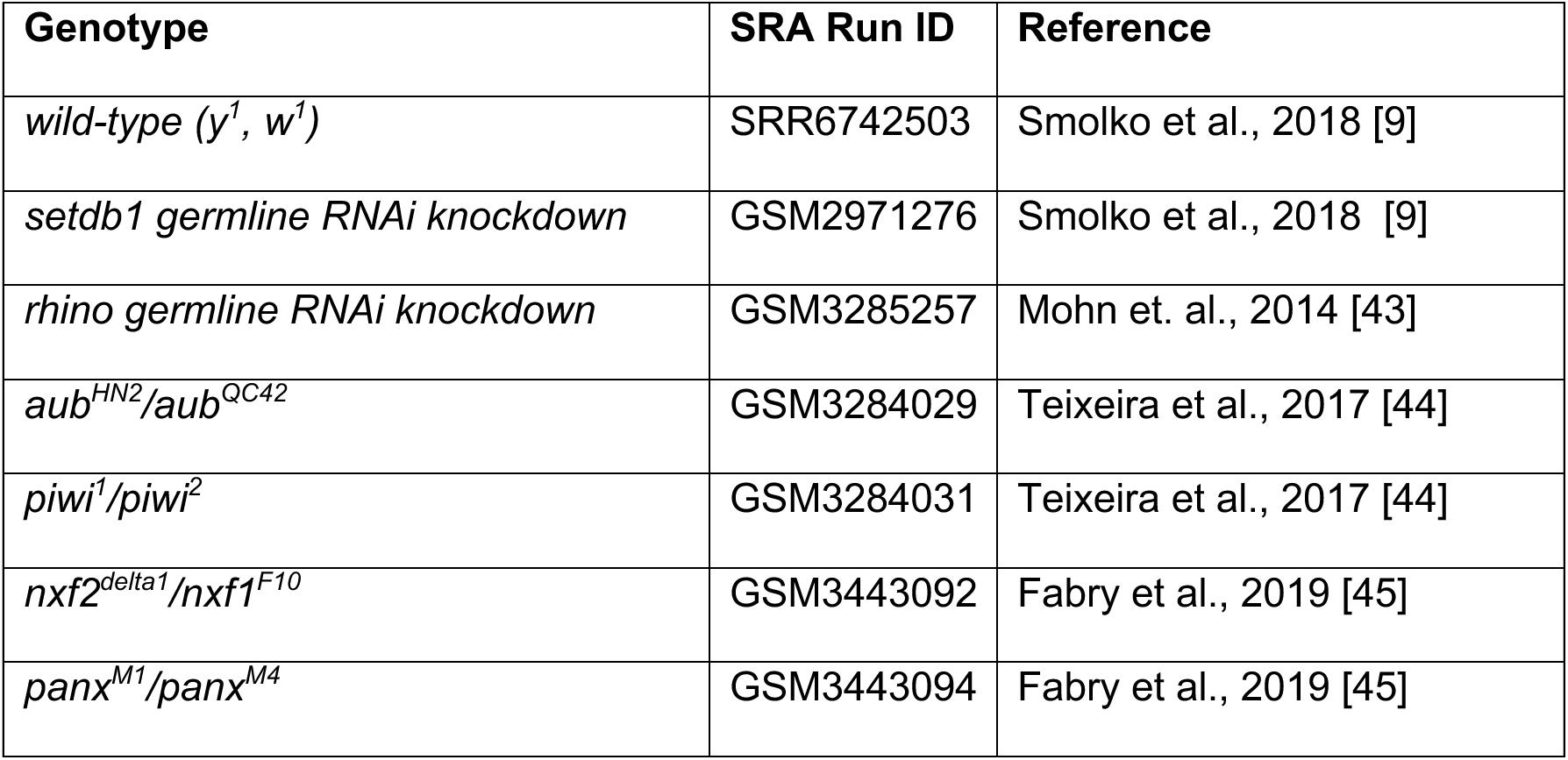
The published RNA-seq data sets utilized in Figure 4.

**Table S4.**
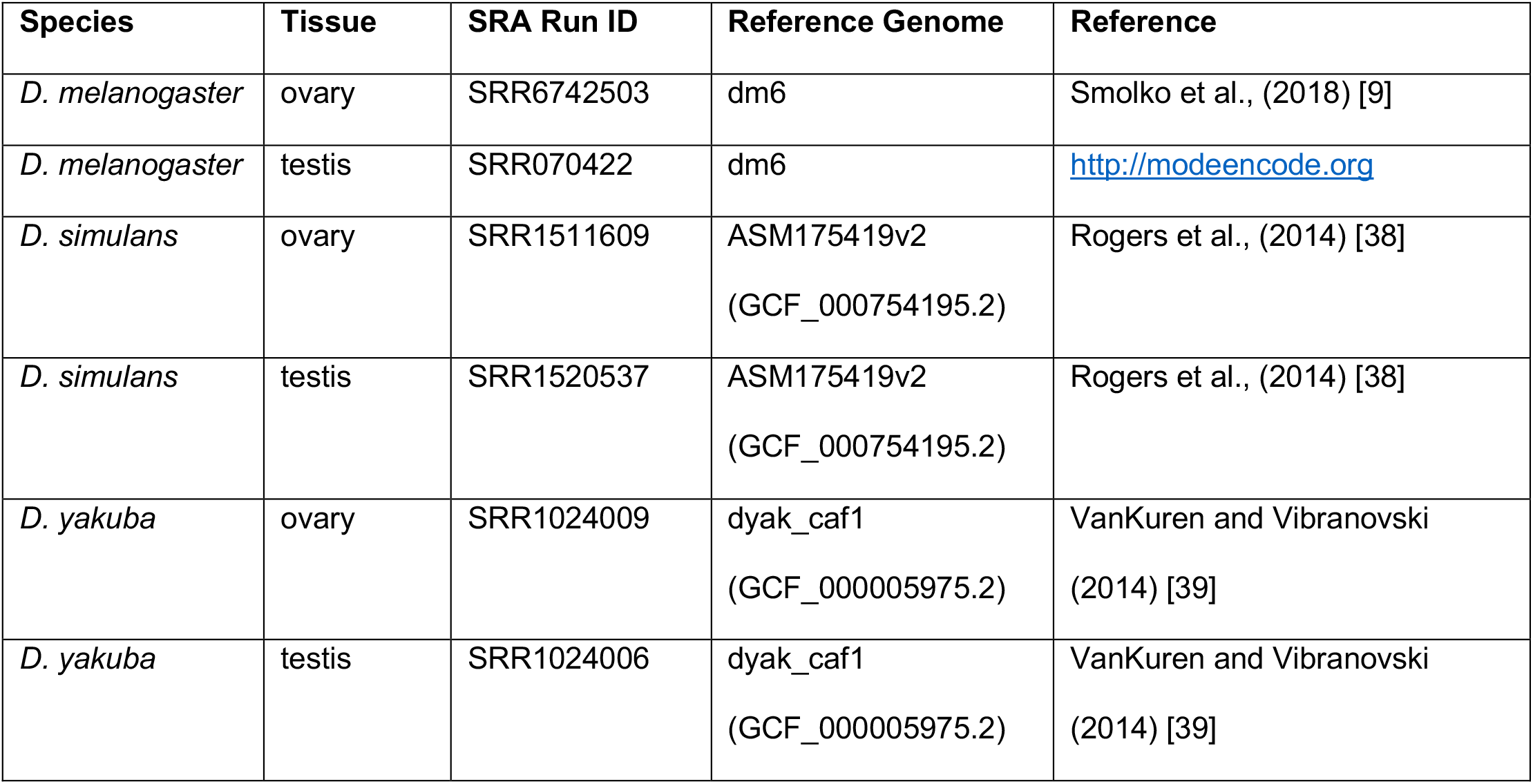
The published RNA-seq data sets utilized in Figure 3.

## Notes

### Competing Interest Statement

The authors have declared no competing interest.

### Summary of Updates

This version of the manuscript has been revised to include additional data and author.

## References

1. Becker JS, Nicetto D, Zaret KS. H3K9me3-Dependent Heterochromatin: Barrier to Cell Fate Changes. Trends in genetics : TIG. 2016;32: 29–41. doi:10.1016/j.tig.2015.11.001

2. Nicetto D, Zaret KS. Role of H3K9me3 heterochromatin in cell identity establishment and maintenance. Current Opinion in Genetics & Development. 2019;55: 1–10. doi:10.1016/j.gde.2019.04.013

3. Ninova M, Tóth KF, Aravin AA. The control of gene expression and cell identity by H3K9 trimethylation. Development (Cambridge, England). 2019;146. doi:10.1242/dev.181180

4. Padeken J, Methot SP, Gasser SM. Establishment of H3K9-methylated heterochromatin and its functions in tissue differentiation and maintenance. Nat Rev Mol Cell Bio. 2022; 1–18. doi:10.1038/s41580-022-00483-w

5. Martienssen R, Moazed D. RNAi and Heterochromatin Assembly. Csh Perspect Biol. 2015;7: a019323. doi:10.1101/cshperspect.a019323

6. Zofall M, Yamanaka S, Reyes-Turcu FE, Zhang K, Rubin C, Grewal SIS. RNA elimination machinery targeting meiotic mRNAs promotes facultative heterochromatin formation. Science. 2012;335: 96–100. doi:10.1126/science.1211651

7. Rechtsteiner A, Costello ME, Egelhofer TA, Garrigues JM, Strome S, Petrella LN. Repression of Germline Genes in Caenorhabditis elegans Somatic Tissues by H3K9 Dimethylation of Their Promoters. Genetics. 2019;212: 125–140. doi:10.1534/genetics.118.301878

8. Methot SP, Padeken J, Brancati G, Zeller P, Delaney CE, Gaidatzis D, et al. H3K9me selectively blocks transcription factor activity and ensures differentiated tissue integrity. Nat Cell Biol. 2021;23: 1163–1175. doi:10.1038/s41556-021-00776-w

9. Smolko AE, Shapiro-Kulnane L, Salz HK. The H3K9 methyltransferase SETDB1 maintains female identity in Drosophila germ cells. Nature communications. 2018;9: 4155. doi:10.1038/s41467-018-06697-x

10. Mochizuki K, Sharif J, Shirane K, Uranishi K, Bogutz AB, Janssen SM, et al. Repression of germline genes by PRC1.6 and SETDB1 in the early embryo precedes DNA methylation-mediated silencing. Nat Commun. 2021;12: 7020. doi:10.1038/s41467-021-27345-x

11. Karimi MM, Goyal P, Maksakova IA, Bilenky M, Leung D, Tang JX, et al. DNA Methylation and SETDB1/H3K9me3 Regulate Predominantly Distinct Sets of Genes, Retroelements, and Chimeric Transcripts in mESCs. Cell Stem Cell. 2011;8: 676–687. doi:10.1016/j.stem.2011.04.004

12. Wu K, Liu H, Wang Y, He J, Xu S, Chen Y, et al. SETDB1-Mediated Cell Fate Transition between 2C-Like and Pluripotent States. Cell Reports. 2020;30: 25-36.e6. doi:10.1016/j.celrep.2019.12.010

13. Jiang Y, Loh Y-HE, Rajarajan P, Hirayama T, Liao W, Kassim BS, et al. The methyltransferase SETDB1 regulates a large neuron-specific topological chromatin domain. Nature genetics. 2017;49: 1239–1250. doi:10.1038/ng.3906

14. Du D, Katsuno Y, Meyer D, Budi EH, Chen S-H, Koeppen H, et al. Smad3-mediated recruitment of the methyltransferase SETDB1/ESET controls Snail1 expression and epithelial-mesenchymal transition. EMBO reports. 2018;19: 135–155. doi:10.15252/embr.201744250

15. Koide S, Oshima M, Takubo K, Yamazaki S, Nitta E, Saraya A, et al. Setdb1 maintains hematopoietic stem and progenitor cells by restricting the ectopic activation of nonhematopoietic genes. Blood. 2016;128: 638–649. doi:10.1182/blood-2016-01-694810

16. Schultz DC, Ayyanathan K, Negorev D, Maul GG, Rauscher FJ. SETDB1: a novel KAP-1-associated histone H3, lysine 9-specific methyltransferase that contributes to HP1-mediated silencing of euchromatic genes by KRAB zinc-finger proteins. Genes & Development. 2002;16: 919–932. doi:10.1101/gad.973302

17. Tan S-L, Nishi M, Ohtsuka T, Matsui T, Takemoto K, Kamio-Miura A, et al. Essential roles of the histone methyltransferase ESET in the epigenetic control of neural progenitor cells during development. Development (Cambridge, England). 2012;139: 3806–3816. doi:10.1242/dev.082198

18. Cheng E-C, Hsieh C-L, Liu N, Wang J, Zhong M, Chen T, et al. The Essential Function of SETDB1 in Homologous Chromosome Pairing and Synapsis during Meiosis. Cell Reports. 2021;34: 108575. doi:10.1016/j.celrep.2020.108575

19. Lohmann F, Loureiro J, Su H, Fang Q, Lei H, Lewis T, et al. KMT1E mediated H3K9 methylation is required for the maintenance of embryonic stem cells by repressing trophectoderm differentiation. Stem cells (Dayton, Ohio). 2010;28: 201–212. doi:10.1002/stem.278

20. Bilodeau S, Kagey MH, Frampton GM, Rahl PB, Young RA. SetDB1 contributes to repression of genes encoding developmental regulators and maintenance of ES cell state. Genes & Development. 2009;23: 2484–2489. doi:10.1101/gad.1837309

21. McCarthy RL, Kaeding KE, Keller SH, Zhong Y, Xu L, Hsieh A, et al. Diverse heterochromatin-associated proteins repress distinct classes of genes and repetitive elements. Nat Cell Biol. 2021;23: 905–914. doi:10.1038/s41556-021-00725-7

22. Nicetto D, Donahue G, Jain T, Peng T, Sidoli S, Sheng L, et al. H3K9me3-heterochromatin loss at protein-coding genes enables developmental lineage specification. Science. 2019;363: 294–297. doi:10.1126/science.aau0583

23. Biferali B, Bianconi V, Perez DF, Kronawitter SP, Marullo F, Maggio R, et al. Prdm16-mediated H3K9 methylation controls fibro-adipogenic progenitors identity during skeletal muscle repair. Sci Adv. 2021;7: eabd9371. doi:10.1126/sciadv.abd9371

24. Marsano RM, Dimitri P. Constitutive Heterochromatin in Eukaryotic Genomes: A Mine of Transposable Elements. Cells. 2022;11: 761. doi:10.3390/cells11050761

25. Wallrath LL, Rodriguez-Tirado F, Geyer PK. Shining Light on the Dark Side of the Genome. Cells. 2022;11: 330. doi:10.3390/cells11030330

26. Janssen A, Colmenares SU, Karpen GH. Heterochromatin: Guardian of the Genome. Annual review of cell and developmental biology. 2018;34: 265–288. doi:10.1146/annurev-cellbio-100617-062653

27. Aravin AA. Pachytene piRNAs as beneficial regulators or a defense system gone rogue. Nat Genet. 2020;52: 644–645. doi:10.1038/s41588-020-0656-8

28. Yang P, Wang Y, Macfarlan TS. The Role of KRAB-ZFPs in Transposable Element Repression and Mammalian Evolution. Trends in genetics : TIG. 2017;33: 871–881. doi:10.1016/j.tig.2017.08.006

29. Ecco G, Imbeault M, Trono D. KRAB zinc finger proteins. Development (Cambridge, England). 2017;144: 2719–2729. doi:10.1242/dev.132605

30. Bruno M, Mahgoub M, Macfarlan TS. The Arms Race Between KRAB–Zinc Finger Proteins and Endogenous Retroelements and Its Impact on Mammals. Annu Rev Genet. 2019;53: 1–24. doi:10.1146/annurev-genet-112618-043717

31. Senft AD, Macfarlan TS. Transposable elements shape the evolution of mammalian development. Nat Rev Genet. 2021;22: 691–711. doi:10.1038/s41576-021-00385-1

32. Benner L, Castro EA, Whitworth C, Venken KJT, Yang H, Fang J, et al. Drosophila Heterochromatin Stabilization Requires the Zinc-Finger Protein Small Ovary. Genetics. 2019;213: 877–895. doi:10.1534/genetics.119.302590

33. Jankovics F, Bence M, Sinka R, Faragó A, Bodai L, Pettkó-Szandtner A, et al. Drosophila small ovary gene is required for transposon silencing and heterochromatin organization, and ensures germline stem cell maintenance and differentiation. Development (Cambridge, England). 2018;145: dev170639. doi:10.1242/dev.170639

34. Andreev VI, Yu C, Wang J, Schnabl J, Tirian L, Gehre M, et al. Panoramix SUMOylation on chromatin connects the piRNA pathway to the cellular heterochromatin machinery. Nat Struct Mol Biol. 2022;29: 130–142. doi:10.1038/s41594-022-00721-x

35. Baumgartner L, Handler D, Platzer SW, Yu C, Duchek P, Brennecke J. The Drosophila ZAD zinc finger protein Kipferl guides Rhino to piRNA clusters. Elife. 2022;11. doi:10.7554/elife.80067

36. Smolko AE, Shapiro-Kulnane L, Salz HK. An autoregulatory switch in sex-specific phf7 transcription causes loss of sexual identity and tumors in the Drosophila female germline. Development (Cambridge, England). 2020. doi:10.1242/dev.192856

37. Yang SY, Baxter EM, Doren M van. Phf7 controls male sex determination in the Drosophila germline. Developmental Cell. 2012;22: 1041–1051. doi:10.1016/j.devcel.2012.04.013

38. Rogers RL, Shao L, Sanjak JS, Andolfatto P, Thornton KR. Revised Annotations, Sex-Biased Expression, and Lineage-Specific Genes in the Drosophila melanogaster Group. G3 Genes Genomes Genetics. 2014;4: 2345–2351. doi:10.1534/g3.114.013532

39. VanKuren NW, Vibranovski MD. A Novel Dataset for Identifying Sex-Biased Genes in Drosophila. J Genom. 2014;2: 64–67. doi:10.7150/jgen.7955

40. Osumi K, Sato K, Murano K, Siomi H, Siomi MC. Essential roles of Windei and nuclear monoubiquitination of Eggless/SETDB1 in transposon silencing. Embo Rep. 2019;20: e48296. doi:10.15252/embr.201948296

41. Rangan P, Malone CD, Navarro C, Newbold SP, Hayes PS, Sachidanandam R, et al. piRNA production requires heterochromatin formation in Drosophila. Current biology : CB. 2011;21: 1373–1379. doi:10.1016/j.cub.2011.06.057

42. Sienski G, Dönertas D, Brennecke J. Transcriptional silencing of transposons by Piwi and maelstrom and its impact on chromatin state and gene expression. Cell. 2012;151: 964–980. doi:10.1016/j.cell.2012.10.040

43. Mohn F, Sienski G, Handler D, Brennecke J. The rhino-deadlock-cutoff complex licenses noncanonical transcription of dual-strand piRNA clusters in Drosophila. Cell. 2014;157: 1364–1379. doi:10.1016/j.cell.2014.04.031

44. Teixeira FK, Okuniewska M, Malone CD, Coux R-X, Rio DC, Lehmann R. piRNA-mediated regulation of transposon alternative splicing in the soma and germ line. Nature. 2017;8: 272. doi:10.1038/nature25018

45. Fabry MH, Ciabrelli F, Munafò M, Eastwood EL, Kneuss E, Falciatori I, et al. piRNA-guided co-transcriptional silencing coopts nuclear export factors. Elife. 2019;8: e47999. doi:10.7554/elife.47999

46. Théron E, Maupetit-Mehouas S, Pouchin P, Baudet L, Brasset E, Vaury C. The interplay between the Argonaute proteins Piwi and Aub within Drosophila germarium is critical for oogenesis, piRNA biogenesis and TE silencing. Nucleic Acids Res. 2018;46: gky695.. doi:10.1093/nar/gky695

47. Yu Y, Gu J, Jin Y, Luo Y, Preall JB, Ma J, et al. Panoramix enforces piRNA-dependent cotranscriptional silencing. Science. 2015;350: 339–342. doi:10.1126/science.aab0700

48. Murano K, Iwasaki YW, Ishizu H, Mashiko A, Shibuya A, Kondo S, et al. Nuclear RNA export factor variant initiates piRNA-guided co-transcriptional silencing. Embo J. 2019;38: e102870. doi:10.15252/embj.2019102870

49. Sienski G, Batki J, Senti K-A, Dönertas D, Tirian L, Meixner K, et al. Silencio/CG9754 connects the Piwi-piRNA complex to the cellular heterochromatin machinery. Genes & Development. 2015;29: 2258–2271. doi:10.1101/gad.271908.115

50. Batki J, Schnabl J, Wang J, Handler D, Andreev VI, Stieger CE, et al. The nascent RNA binding complex SFiNX licenses piRNA-guided heterochromatin formation. Nat Struct Mol Biol. 2019;26: 720–731. doi:10.1038/s41594-019-0270-6

51. Zhao K, Cheng S, Miao N, Xu P, Lu X, Zhang Y, et al. A Pandas complex adapted for piRNA-guided transcriptional silencing and heterochromatin formation. Nat Cell Biol. 2019;21: 1261–1272. doi:10.1038/s41556-019-0396-0

52. Chung H-R, Schäfer U, Jäckle H, Böhm S. Genomic expansion and clustering of ZAD-containing C2H2 zinc-finger genes in Drosophila. EMBO reports. 2002;3: 1158–1162. doi:10.1093/embo-reports/kvf243

53. Chung H-R, Löhr U, Jäckle H. Lineage-specific expansion of the zinc finger associated domain ZAD. Molecular biology and evolution. 2007;24: 1934–1943. doi:10.1093/molbev/msm121

54. Kasinathan B, Colmenares SU, McConnell H, Young JM, Karpen GH, Malik HS. Innovation of heterochromatin functions drives rapid evolution of essential ZAD-ZNF genes in Drosophila. eLife. 2020;9. doi:10.7554/elife.63368

55. Shapiro-Kulnane L, Bautista O, Salz HK. An RNA-interference screen in Drosophila to identify ZAD-containing C2H2 zinc finger genes that function in female germ cells. G3 Genes Genomes Genetics. 2020;11: jkaa016. doi:10.1093/g3journal/jkaa016

56. Jauch R, Bourenkov GP, Chung H-R, Urlaub H, Reidt U, Jäckle H, et al. The zinc finger-associated domain of the Drosophila transcription factor grauzone is a novel zinc-coordinating protein-protein interaction module. Structure (London, England : 1993). 2003;11: 1393–1402. doi:10.1016/j.str.2003.09.015

57. Zolotarev N, Fedotova A, Kyrchanova O, Bonchuk A, Penin AA, Lando AS, et al. Architectural proteins Pita, Zw5,and ZIPIC contain homodimerization domain and support specific long-range interactions in Drosophila. Nucleic Acids Res. 2016;44: 7228–7241. doi:10.1093/nar/gkw371

58. Maksimenko O, Kyrchanova O, Klimenko N, Zolotarev N, Elizarova A, Bonchuk A, et al. Small Drosophila zinc finger C2H2 protein with an N-terminal zinc finger-associated domain demonstrates the architecture functions. Biochimica Et Biophysica Acta Bba - Gene Regul Mech. 2020;1863: 194446. doi:10.1016/j.bbagrm.2019.194446

59. Bonchuk AN, Boyko KM, Nikolaeva AY, Burtseva AD, Popov VO, Georgiev PG. Structural insights into highly similar spatial organization of zinc-finger associated domains with a very low sequence similarity. Structure. 2022;30: 1004-1015.e4. doi:10.1016/j.str.2022.04.009

60. Bonchuk A, Boyko K, Fedotova A, Nikolaeva A, Lushchekina S, Khrustaleva A, et al. Structural basis of diversity and homodimerization specificity of zinc-finger-associated domains in Drosophila. Nucleic Acids Res. 2021;49: 2375–2389. doi:10.1093/nar/gkab061

61. Shapiro-Kulnane L, Smolko AE, Salz HK. Maintenance of Drosophila germline stem cell sexual identity in oogenesis and tumorigenesis. Development (Cambridge, England). 2015;142: 1073–1082. doi:10.1242/dev.116590

62. Hinnant TD, Merkle JA, Ables ET. Coordinating Proliferation, Polarity, and Cell Fate in the Drosophila Female Germline. Frontiers in cell and developmental biology. 2020;8: 19. doi:10.3389/fcell.2020.00019

63. Chau J, Kulnane LS, Salz HK. Sex-lethal facilitates the transition from germline stem cell to committed daughter cell in the Drosophila ovary. Genetics. 2009;182: 121–132. doi:10.1534/genetics.109.100693

64. Chau J, Kulnane LS, Salz HK. Sex-lethal enables germline stem cell differentiation by down-regulating Nanos protein levels during Drosophila oogenesis. Proceedings of the National Academy of Sciences of the United States of America. 2012;109: 9465–9470. doi:10.1073/pnas.1120473109

65. Li Y, Minor NT, Park JK, Mckearin DM, Maines JZ. Bam and Bgcn antagonize Nanos-dependent germ-line stem cell maintenance. Proceedings of the National Academy of Sciences. 2009;106: 9304–9309. doi:10.1073/pnas.0901452106

66. Yi X, Vries HI de, Siudeja K, Rana A, Lemstra W, Brunsting JF, et al. Stwl modifies chromatin compaction and is required to maintain DNA integrity in the presence of perturbed DNA replication. Molecular biology of the cell. 2009;20: 983–994. doi:10.1091/mbc.e08-06-0639

67. Shokri L, Inukai S, Hafner A, Weinand K, Hens K, Vedenko A, et al. A Comprehensive Drosophila melanogaster Transcription Factor Interactome. Cell Reports. 2019;27: 955-970.e7. doi:10.1016/j.celrep.2019.03.071

68. Zinshteyn D, Barbash DA. Stonewall prevents expression of ectopic genes in the ovary and accumulates at insulator elements in D. melanogaster. Plos Genet. 2022;18: e1010110. doi:10.1371/journal.pgen.1010110

69. Allshire RC, Madhani HD. Ten principles of heterochromatin formation and function. Nature reviews Molecular cell biology. 2018;19: 229–244. doi:10.1038/nrm.2017.119

70. Sarkar K, Kotb NM, Lemus A, Martin ET, McCarthy A, Camacho J, et al. A feedback loop between heterochromatin and the nucleopore complex controls germ-cell to oocyte transition during Drosophila oogenesis. Biorxiv. 2021; 2021.10.31.466575. doi:10.1101/2021.10.31.466575

71. Sarov M, Barz C, Jambor H, Hein MY, Schmied C, Suchold D, et al. A genome-wide resource for the analysis of protein localisation in Drosophila. Elife. 2016;5: e12068. doi:10.7554/elife.12068

72. Hu Y, Comjean A, Rodiger J, Liu Y, Gao Y, Chung V, et al. FlyRNAi.org—the database of the Drosophila RNAi screening center and transgenic RNAi project: 2021 update. Nucleic Acids Res. 2020;49: gkaa936.. doi:10.1093/nar/gkaa936

73. Gratz SJ, Ukken FP, Rubinstein CD, Thiede G, Donohue LK, Cummings AM, et al. Highly specific and efficient CRISPR/Cas9-catalyzed homology-directed repair in Drosophila. Genetics. 2014;196: 961–971. doi:10.1534/genetics.113.160713

74. Port F, Strein C, Stricker M, Rauscher B, Heigwer F, Zhou J, et al. A large-scale resource for tissue-specific CRISPR mutagenesis in Drosophila. Elife. 2020;9: e53865. doi:10.7554/elife.53865

75. Livak KJ, Schmittgen TD. Analysis of relative gene expression data using real-time quantitative PCR and the 2(-Delta Delta C(T)) Method. Methods (San Diego, Calif). 2001;25: 402–408. doi:10.1006/meth.2001.1262

